# Seasonal contact and migration structure mass epidemics and inform outbreak preparedness in bottlenose dolphins

**DOI:** 10.1101/2024.10.02.616317

**Authors:** Melissa Collier, Kim Urian, Sarah Theisen, Ann-Marie Jacoby, Sarah Wilkin, Eric M. Patterson, Megan M. Wallen, Vittoria Colizza, Janet Mann, Shweta Bansal

**Affiliations:** Department of Biology, Georgetown University, Washington DC; Marine Science and Conservation Division, Duke University Marine Lab, Beaufort, North Carolina; Office of Protected Resources, NOAA Fisheries, Silver Spring, MD, USA; Protected Resources Division, West Coast Region, NOAA Fisheries, Seattle, WA, USA; Sorbonne Université, INSERM, Institut Pierre Louis d’Épidémiologie et de Santé Publique, Paris, France; Department of Psychology, Georgetown University, Washington DC

## Abstract

Infectious respiratory diseases have detrimental impacts across wildlife taxa, particularly in marine species. Despite this vulnerability, we lack information on the complex spatial and contact structures of marine populations which reduces our ability to understand disease spread and our preparedness for epidemic response. We leveraged a collated dataset to establish the first data-driven epidemiological model on a cetacean species, the Tamanend’s bottlenose dolphin (*Tursiops erebennus*), whose populations are periodically impacted by deadly respiratory disease in the northwest Atlantic. We found their spatial distribution and contact is heterogeneous along the coastline and varies by ecotype, which explains differences in infection burdens observed in past outbreaks. We also determined that outbreaks beginning in northern parts of their habitat during migratory seasons have the highest epidemic risk and that dolphins in North Carolina estuaries would be the best sentinels for disease surveillance. Our mathematical model provides a generalizable, non-invasive tool that takes advantage of routinely collected marine mammal data to mechanistically understand disease transmission and inform disease surveillance tactics for marine sentinels. Our findings highlight the heterogeneities that play a crucial role in shaping the impacts of infectious diseases in wildlife, and how a data-driven understanding of these mechanisms can enhance epidemic preparedness.

## 1.0 Introduction

Infectious diseases have detrimental impacts on wildlife^1,2^, but they remain understudied in marine populations despite their significant impact on these ocean-dwelling species. The effect of disease on cetaceans (whales, porpoises, and dolphins) is of particular interest, as they are predators and can serve as sentinel species; their protection is vital for maintaining a balanced and healthy ocean ecosystem^3–5^, and their populations are currently under threat of intensified pathogen outbreaks due to the direct and indirect effects of climate change^6^. However, few data exist on the spatial and social structures of these species^7,8^, consequently limiting our ability to analyze, forecast, and respond to disease threats. To address this, we leverage multiple individual-level datasets on the behavior of a sentinel cetacean species that is periodically impacted by deadly respiratory disease. We develop a novel mathematical modeling approach to 1) characterize how sociality and space use impact the structure of vulnerability in a complex marine population, and 2) inform preparedness for future deadly outbreaks in this vulnerable and understudied taxa.

Respiratory transmitted pathogens are responsible for disease related population declines in wildlife including bighorn sheep^9^, mountain goats^10^, house finches^11^, lizards^12^, and tortoises^13^, since they spread quickly via respiratory secretions over both direct and indirect transmission routes^14^. Pathogens of the family morbillivirus are particularly prevalent and deadly in the marine environment^15,16^ with dolphin morbillivirus (DMV) being the most devastating among dolphins^16^. For instance, the virus killed more than 1000 striped dolphins (*Stenella coeruleoalba*) in the coastal waters of Spain, Italy and France in the 1990s^17^, with additional viral resurgences in 2007 and 2011^18,19^. But perhaps the most detrimental DMV outbreaks on record occurred in Tamanend’s bottlenose dolphins (*Tursiops erebennus*, formerly classified as *Tursiops truncatus*^20^) along the Atlantic coast of the United States which depleted some populations by more than 50% in 1987^21^ and 2013^22^, both declared “Unusual Mortality Events”. To analyze respiratory disease spread, forecast outbreaks, or develop surveillance programs for vulnerable marine species like Tamanend’s dolphins, it’s crucial to understand the spatial distribution and contact structure that drive respiratory pathogen transmission in wild animal populations.

It is well known that animal movement such as dispersal (short-distance regular movement to different habitats) and migration (long-distance repeated movements on a seasonal basis) enhances the spatial spread of pathogens both within and across species^7^. Increased effort in collecting marine mammal stranding data has improved our understanding of the spatial distribution of respiratory disease in the marine environment^15,16,23^. Because pathogens rely on host movement to disperse, the spatiotemporal behavior of a host population will determine their risk of exposure and transmission^24^. However, without spatiotemporal data on the associated hosts, pathogen distribution and spread is challenging to predict. For example, Tamanend’s dolphins comprise multiple populations between the New York and Florida coastlines that overlap in space and time^25^, the degree to which is still not well understood. While research along the Georgia^26^ and Florida^27^ coastlines shows seasonal shifts in distribution and spatial overlap between populations of Tamanend’s dolphins, these studies are limited to specific near-shore regions and are unable to make inferences across their habitat range. These constraints reduce our ability to identify or predict the distribution and spread of future outbreaks of respiratory disease in this species.

While host space use controls the spatial distribution of pathogens, host contact controls transmission within populations. Thus host social systems^8^ and individual heterogeneities in contact behavior^28,29^ will dictate disease dynamics. Cetaceans exhibit a variety of social structures that range from being relatively solitary (e.g. baleen whales), to highly gregarious (e.g. dolphins)^30^ suggesting subsequent variability in how disease transmits in this taxa. Tamanend’s dolphins are an extremely social species^31^ that use a behavior known as synchronized breathing (two or more individuals surfacing and breathing simultaneously and in close proximity^32^) to build lifelong social bonds^33–35^, which subsequently puts them at high risk for the transmission of respiratory pathogens^8,16^. The structure of synchronized breathing contact had been virtually unknown in Tamanend’s dolphins, until recent work showed that it differs across age and sex classes with potential respiratory infection consequences^36^. However, our ability to predict seasonal variation in disease risk due to social behavior remains limited.

Despite motivation from past epidemics to study Tamanend’s dolphins’ spatial distribution and contact, the complexity of these processes has hindered inferences about their impact on disease spread in this species as a whole. Tamanend’s dolphins are managed as “stocks” by the National Oceanic and Atmospheric Administration (NOAA). These stocks are distributed along the U.S. eastern seaboard from New York to Florida^25^ and make up two main ecotypes: “estuarine” stocks which primarily occupy bays, rivers, and tributaries closer to shore and engage in small range spatial dispersion, and “coastal” stocks which are found mostly along ocean coastlines, deeper water, and often exhibit larger migratory movements^25^. Historically, the burden of DMV in this species has not been distributed homogeneously along the coastline or by ecotype. For example, during the past two DMV outbreaks the majority of detected infections occurred at northern latitudes in warmer months, while at lower latitudes infections peaked much later and with a lower burden^37,38^ (Figure 1). Further, DMV has clearly depleted coastal stocks^21,39^, while estuarine stocks were seemingly less impacted^40^. Only one study has analyzed the spatial spread of DMV in Tamanend’s dolphins along the US coastline, to infer key infection parameters like its basic reproduction number (R0) and infectious period. However, a significant gap persists as no mechanistic approach has yet been developed to capture spatial heterogeneity and inform ecotype-specific surveillance and preparedness in systems such as these, where typical disease intervention approaches are not logistically or legally feasible.

**Figure 1:**
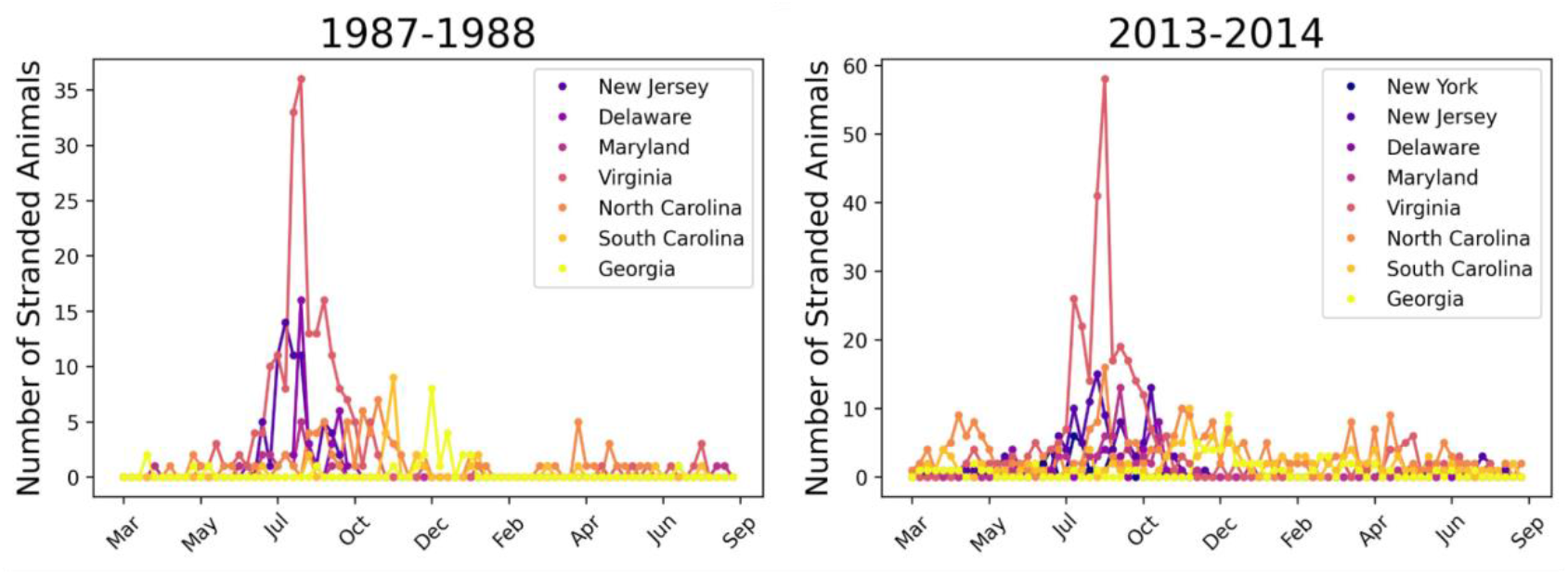
Bottlenose dolphin strandings detected during two DMV outbreaks along the US Atlantic coastline. During both the epidemics beginning in 1987 (left) and 2013 (right), confirmed strandings (when a sick, injured or dead dolphin is found floating or washed ashore) of bottlenose dolphins peaked in July and August along the northern part of the US Atlantic coastline, with smaller peaks at the southern part of the coastline in October-December. While there is notable spatial heterogeneity in strandings along the coastline during both outbreaks, the spatiotemporal distribution of infections was remarkably similar between the two epidemics.

Given their complex socio-spatial dynamics, epidemiological models driven by movement and contact behaviors collected along the entire habitat range of bottlenose dolphins are needed. Metapopulation models provide an approach to model disease dynamics in spatially separated but connected populations for answering diverse epidemiological questions related to pathogen invasion dynamics, persistence, control measures, and evolution^41^. Successful models applied in wildlife systems have described the persistence of lyssavirus^42^ and spread of white-nose syndrome^43^ in bat species, the spread of bovine tuberculosis amongst the migratory brushtail possum^44^, and the prevalence of sarcoptic mange and canine distemper in Yellowstone wolves^45^. Only one study has ever applied metapopulation inference to disease spread among marine mammals, to determine persistence thresholds for respiratory viruses in harbor seals^46^, with none applied in cetaceans.

We developed the first data-driven epidemiological metapopulation model in a cetacean population to model respiratory disease dynamics in Tamanend’s bottlenose dolphins along the Atlantic coast from New York to Florida. To parameterize this model and overcome past limitations, we 1) leveraged a collated dataset containing bottlenose dolphin sightings across 28 different field locations along the Atlantic coastline from New Jersey to Georgia in the US and collected individual-level data at one of these locations on synchronized breathing contact for individuals of both coastal and estuarine ecotypes. We validated this model with outbreak data from the 1987 and 2013 DMV epidemics in this species, and used it to inform our understanding of the role of seasonality in environmental factors and social behavior on disease dynamics. Finally, we applied our best model to assess epidemic risk along the Tamanend’s dolphin habitat range, and identify potential sites for sentinel DMV surveillance. With this work, we characterize the impact of metapopulation structure on disease risk and demonstrate our model’s applicability in improving our preparedness for future outbreaks in vulnerable populations.

## 2.0 Results

We constructed an epidemiological model to assess the impact of metapopulation structure on the dynamics of past outbreaks (Figure 1). First, we inferred metapopulation structure of Tamanend’s dolphins using sighting data from the Mid-Atlantic Bottlenose Dolphin Catalog (MABDC) obtained on dolphins of coastal and estuarine ecotypes between New Jersey and Georgia, and behavioral data from the Potomac-Chesapeake Dolphin Project (PCDP) obtained on both ecotypes from the Chesapeake Bay, where the plurality of DMV infections occurred in 2013. Due to a lack of consistent sighting data contributions from Florida field sites to the MABDC, we did not consider this part of the coastline in our primary analysis, but see Supplementary Information section 10 for an analysis including this region. Given the diverse movement patterns exhibited by Tamanend’s dolphin ecotypes^25^, we use the broader term ‘movement’ when referring to rates of estuarine dispersal and coastal migration.

We used MABDC data to divide the United States Atlantic coastline into ecologically relevant metapopulation “patches”, and estimate movement rates of ecotypes among these patches; we then used PCDP data to estimate the contact driven DMV transmission rates for ecotypes within these patches (Figure 2). To compare our model predictions to past DMV epidemic dynamics, we used data available from the 1987 and 2013 DMV outbreaks in Tamanend’s dolphins obtained from the Smithsonian Division of Marine Mammal Collections and NOAA, respectively. We also estimated the infection burden in coastal and estuarine ecotypes to evaluate whether our models align with expert opinion that coastal ecotypes experienced a higher infection burden^22,25,47^. Finally, we applied our model to inform our preparedness to future DMV outbreaks.

**Figure 2:**
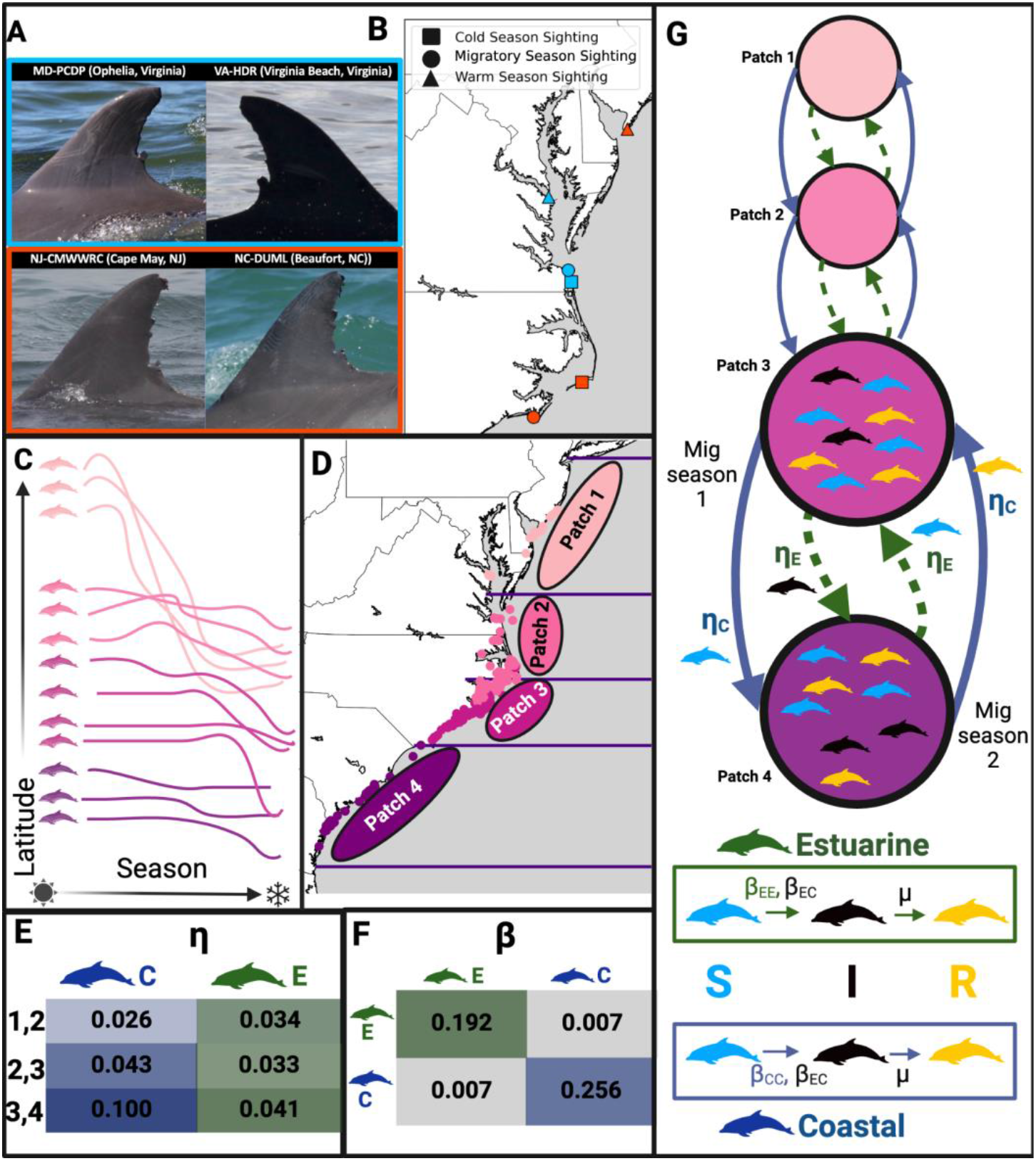
Methods and structure of the epidemiological metapopulation model for the Tamanend’s bottlenose dolphins. (A) First we matched photos of dorsal fins across 28 photo-ID catalogs from New Jersey to Georgia that contribute to the MABDC to (B) establish sighting histories for 410 individuals in the warm water, migratory and cold water seasons. (C) These provide a time series of latitudinal locations which we algorithmically group into four clusters. (D) We used the warm-water season sightings of individuals in each resulting cluster (points, colored by cluster) to establish ecologically relevant metapopulation patches (ellipses). (E) The average movement rates (*η*) of individuals in coastal (blue) and estuarine (green) ecotypes between the patches during the migratory seasons are determined by fitting a multistate capture-recapture model to individual sighting histories. (F) The transmission rates (*β*) within and between the two ecotypes are established with synchronized breathing contact field data collected by the PCDP. (G) The resulting epidemiological metapopulation SIR model allows DMV to move between patches based on *η*, and individuals move from susceptible (S) to infected (I) based on *β*, and recover at recovery rate *μ*. Dorsal fin photos (A) courtesy of the PCDP, Cape May Whale Watch and Research Center (Melissa Laurino), Duke University Marine Lab (Kim Urian), and HDR Inc (Amy Engelhaupt), taken under NMFS permit nos: 19403, 808-1798, 16239. Figure created with Biorender.com.

### 2.1 Metapopulation structure can reproduce historical epidemic dynamics

Using empirical sighting and contact data from the MABDC and the PCDP, we produced novel estimates of Tamanend’s bottlenose dolphin 1) metapopulation patches along the Atlantic coast, 2) movement rates between these patches, and 3) contact driven transmission rates within patches. We used these results to model the spread of DMV along the Atlantic coast and compared the resulting infection dynamics to those of past DMV outbreaks.

#### 2.1.1 Dolphin movement and contact varies by ecotype

Using sighting histories established for 410 dolphins with MABDC data (Figure 2A,B) we clustered dolphins based on similar spatiotemporal sightings (Figure 2C). We determined the optimal number clusters is four (Figure S1), and thus divided the coastline into four metapopulation patches based on the average warm water season (defined as July-September, Figure S4) habitat ranges of dolphins in each cluster (Figure 2D), as this time of year is when stocks of dolphins are thought to be most delineated from each other^25^. The resulting patch 1 occupies the coastlines of the northernmost parts of our study area (New Jersey to northern Virginia), followed by patch 2 (southern Virginia to northern North Carolina), patch 3 (southern North Carolina), and patch 4 (South Carolina and Georgia). A comparison to NOAA’s presumed ranges for Tamanend’s dolphin stocks suggests that our patch delineation is valid (Figure S2).

Using MABDC sighting data from dolphins of different ecotypes (coastal n = 37; estuarine n = 373) across 28 field sites along the Atlantic coast, we estimated movement rates of both coastal (*η*_*C*_) and estuarine (*η*_*E*_) individuals between our defined patches. We found that these movement rates are generally higher for coastal individuals compared to estuarine individuals except between patches 1 and 2. For both estuarine and coastal individuals, the highest rate of movement occurs between patches 3 and 4 (Figure 2E, Table S7).

Using behavioral data on coastal (n=16) and estuarine (n=55) individuals from the PCDP, we estimated that mean synchrony degree (i.e., the average number of synchronized breathing contacts capable of transmitting DMV) is *K*_*c*_ =8 individuals per day for coastal individuals and *k*_*E*_=6 individuals per day for estuarine individuals. We also empirically estimated synchrony mixing (i.e., the proportion of contacts of one ecotype that are with the other ecotype) between estuarine and coastal individuals (*α*) to range between 0 and 0.06. Combining synchrony degree, synchrony mixing, and the infectivity of DMV (*τ*= 0.032), we estimated the ecotype-specific transmission rates within (*β*_*CC*_, *β*_*EE*_) and between (*β*_*CE*_, *β*_*EC*_) ecotypes, and predicted transmission rates between coastal individuals (*β*_*CC*_=*0*.*256*) to be higher than those between estuarine individuals (*β*_*EE*_=*0*.*192*); within ecotype transmission rates were predicted to be higher than between ecotype transmission rates (*β*_*CE*_ and *β*_*EC*_ = 0.007) (Figure 2F).

#### 2.1.2 Seasonal changes in contact best explain past infection dynamics

Based on the metapopulation dynamics and transmission rates we empirically estimated, we predicted epidemic dynamics in Tamanend’s dolphins along the Atlantic coast using a metapopulation epidemiological model (Figure 2G). In this model, estuarine movement (i.e., dispersal) occurs year round, and coastal movement (i.e., migration) occurs seasonally. We used least squares fitting and Pearson’s correlations to observed epidemic dynamics, and found that model predictions for infected animals over time are most consistent with data from the 1987 and 2013 outbreaks when the outbreak onset is assumed to be in patch 2 between May 1-May 10 (Figure S8).

Next, we tested hypotheses about the impact of seasonal changes due to environmental factors and social behavior on transmission dynamics. In the first scenario, we considered transmission rates {*β*} to be reduced by 20% outside of the cold water season to represent higher DMV infectiousness in colder environments where enveloped viruses such as DMV are thought to be more effectively transmitted^48^. We find this model is less consistent with the 1987 and 2013 outbreak dynamics (Figure 3C,D,E, S9). In a second scenario, we considered {*β*} to be reduced by 20% outside of April-July to represent a higher breathing synchrony degree during breeding seasons when contact rates are typically higher^49^. We find this scenario better predicts observed dynamics (Figure 3D,E, S9) by reducing infection peaks in patches 1, 3, and 4 (Figure 3B) as dolphins are most prevalent in these patches when *β* is lower, and allowing for a longer epidemic period from the reduced *β* outside the short breeding season. Notably, other realistic changes to breeding and coldwater season lengths and *β* reduction values do not qualitatively affect these results (Table S6). This indicates contact structured by seasonal social behavior may explain the spatial heterogeneity and temporal dynamics of DMV outbreaks.

**Figure 3.**
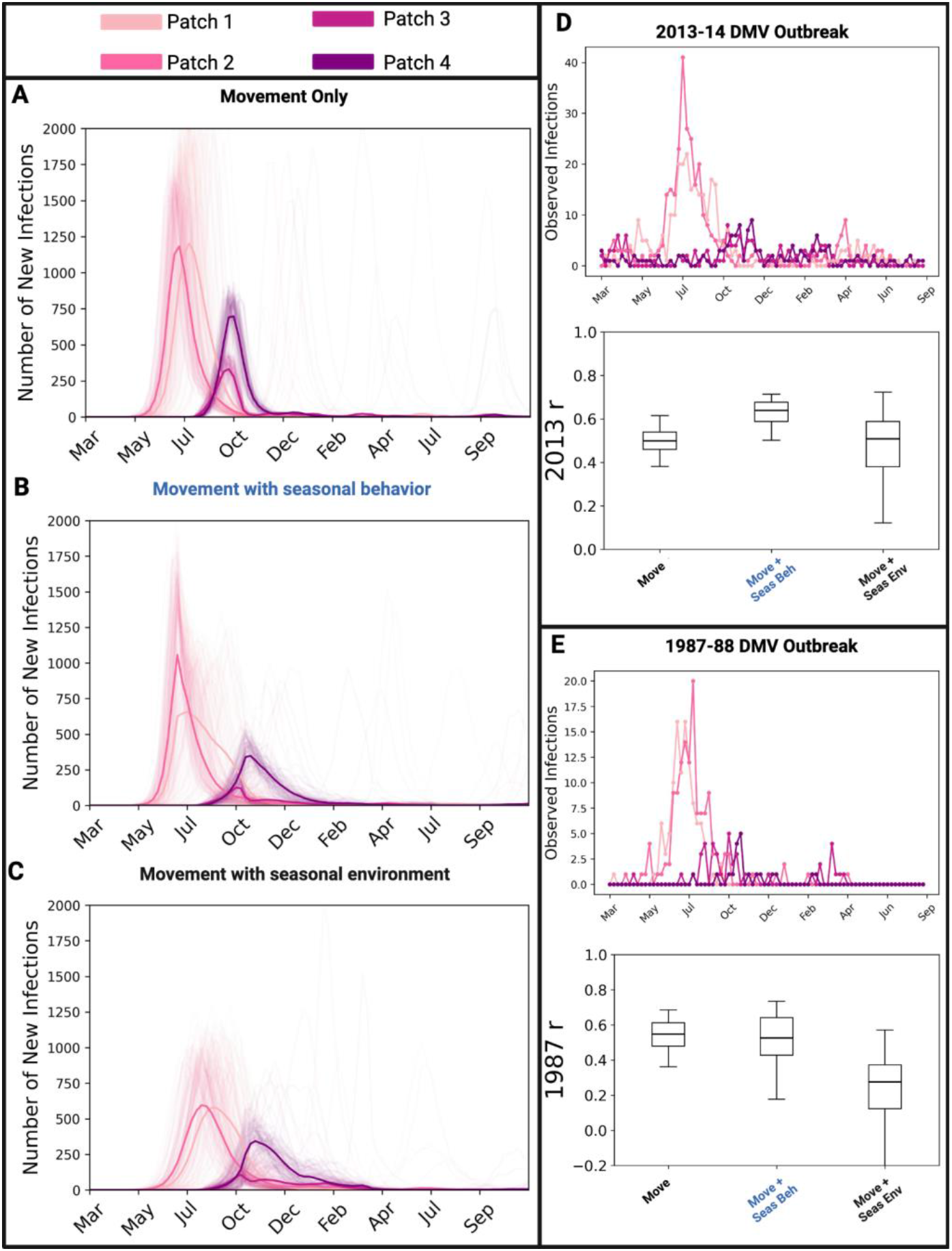
The effect of seasonality in transmission on disease dynamics. The infection time series of a model that considers differences in contact among ecotypes and (A) only dolphin movement with no seasonal changes to transmission rates; (B) dolphin movement and higher transmission rate due to behavior in the breeding season; and (C) dolphin movement with higher transmission due to the environment in the cold water season. We see that a model with contact structured by seasonal behavior (B), indicated in blue, is most consistent with DMV outbreak data from the 2013 (D) and 1987 (E) epidemics based on the Pearson’s correlation coefficients (r) and visual comparisons of the time series. Figure partially created with Biorender.com.

While the Florida coastline was undoubtedly impacted by DMV, there is a lack of consistent, available, sighting data in this region to estimate metapopulation structure in that part of the coastline. To inform what movement to Florida is best supported by the observed disease dynamics, we produced counterfactuals using our epidemiological model. We found the most likely avenue of infection into this area would occur with the migration of coastal individuals into Florida waters from patch 4 (97% epidemic likelihood), as opposed to the sporadic interactions of dolphins from between these two regions (33% epidemic likelihood) (Figure S16).

### 2.2 Movement and contact structure significantly impacts ecotype vulnerability

Best available data suggests that the coastal ecotype was more vulnerable than the estuarine ecotype^22,25,47^ to DMV infection. Our empirically-informed epidemiological metapopulation model predicts higher relative infection burden for coastal individuals compared to estuarine individuals, with a larger bias in burden when seasonal social behavior is incorporated (Figure S10).

To mechanistically understand the impact of contact and movement on ecotype vulnerabilities, we systematically controlled and varied different components of the metapopulation structure in our model. Using the model that was most consistent with observed outbreak dynamics (the seasonal social behavior model), we considered a series of control models in which we remove the variation in 1) ecotype specific movement, 2) overall movement, or 3) ecotype contact.

When movement rates were controlled (i.e., assumed a homogeneous movement rate across patches for each ecotype), the relative infection burden of coastal to estuarine individuals is unaffected (Figure 4, S11), but infection in patch 1 occurred much later than observed in past epidemics (Figure S12). This was also observed to a lesser degree when movement rates were also controlled by ecotype (i.e., assumed the same homogenous movement rate across patches and ecotypes) (Figure S12). When ecotype contact structure was controlled (i.e., assumed the same homogenous transmission rate{*β*}for both ecotypes), the relative disease burden between coastal and estuarine infections was more biased toward estuarine infection (Figure 4). Our findings thus suggest that spatial and ecotype heterogeneity in movement are important for describing infection dynamics at epidemic onset, while ecotype heterogeneity in contact may explain higher coastal ecotype vulnerability to DMV along the coastline.

**Figure 4.**
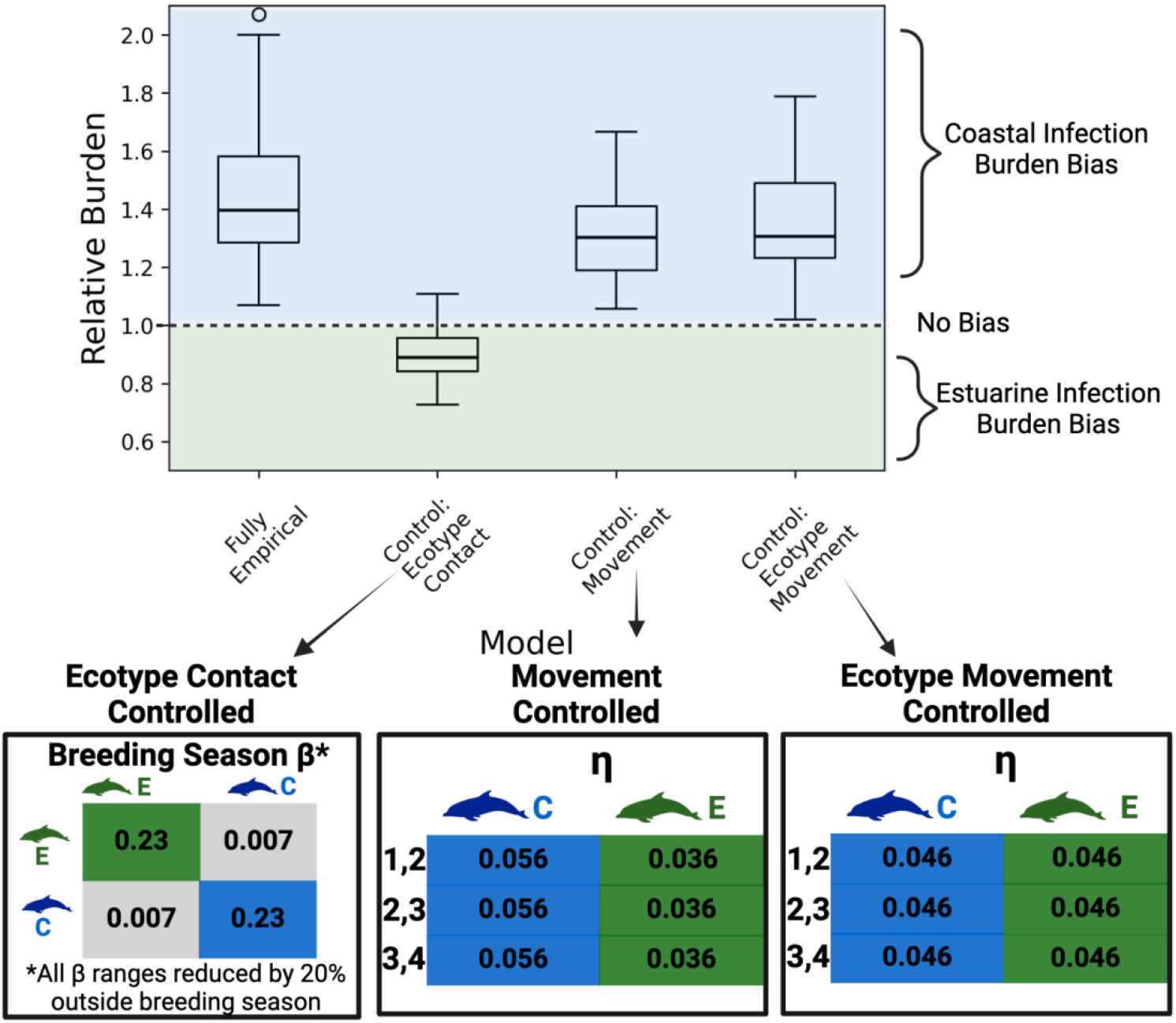
The effect of movement and contact on ecotype infection burden. Relative burden is the ratio of the proportion of coastal individuals infected to estuarine individuals infected. When relative burden is equal to 1 (dotted line), there is no bias in infection burden between the two ecotypes. Values greater than 1 indicate a bias towards coastal infections, and values less than 1 indicate a bias towards estuarine infections. We controlled different components of the metapopulation structure captured in Figure 2 (Fully Empirical) to examine their effect on relative burden by removing heterogeneity in our *β* or *η* estimates. Figure partially created with Biorender.

### 2.3 Disease surveillance and epidemic risk assessment is optimized by leveraging movement and contact structure

To inform outbreak preparedness and optimize surveillance for future DMV outbreaks, we examined how epidemic risk (a combination of the size of the infected population and the likelihood of an epidemic occurring) varies with the onset location and time period, and how particular populations along the coast might serve as sentinels for DMV surveillance.

The epidemic risk of our seasonal social behavior model where infection begins between May 1-10 in patch 2 was 69%. Out of 48 alternative onset location and time scenarios, only seven resulted in similar (60-80%) epidemic risk. Of these similar scenarios, most supported epidemic onset during the migratory seasons in patches 1 or 2 and no scenario with epidemic onset in patch 4 or during the warm or cold water seasons carried the same epidemic risk (Figure 5A). Further, high epidemic risk scenarios with onset in migratory season 1 resulted in epidemic dynamics similar to past DMV outbreaks compared to high risk scenarios with onset in migratory season 2 (Figure 5AB, S14).

**Figure 5.**
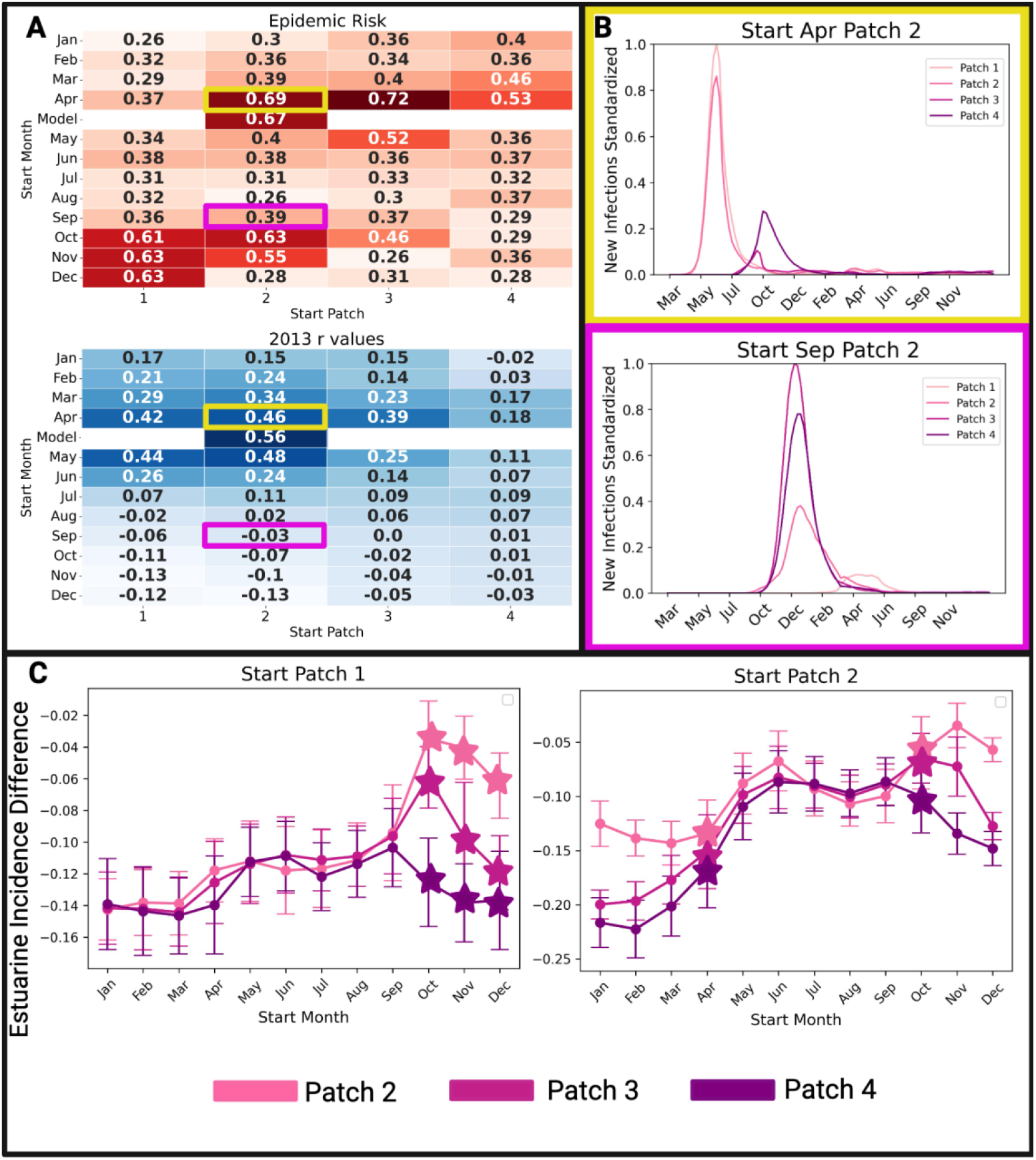
Applying the model to inform DMV preparedness. (A) The epidemic risk values for DMV infection beginning in all possible onset scenarios and their associated Pearson’s correlation coefficients (r) to the 2013 outbreak data. These values show that outbreaks beginning in patches 1 and 2 have the highest epidemic risk but not all of these scenarios result in the dynamics observed in past outbreaks. (B) The average infection time series for two epidemics with high risks show that outbreaks starting in migratory season 1 (yellow) result in dynamics more similar to historic DMV epidemics than outbreaks beginning in migratory season 2 (pink). (C) The difference in the weekly infection incidences of estuarine individuals (the total number of new infections in each patch, each week, normalized by the number of estuarine individuals present) in patches 2, 3 and 4 and that of the entire coastline when outbreaks start in the higher risk patches 1 and 2. Stars indicate scenarios with high epidemic risk (from A). Figure partially created with Biorender.com.

While it is assumed that coastal populations bore a higher infection burden in past DMV outbreaks, estuarine individuals are more accessible for data collection given their smaller, predictable, and nearshore habitat ranges. Thus we examined the possibility of estuarine populations serving as sentinels for DMV monitoring for the entire Atlantic coast. We compared the average weekly infection incidence of estuarine individuals (in patches 2, 3, 4 as these patches currently have established estuarine populations^25^) to the average weekly infection incidence for the entire coastline (coastal + estuarine incidence in all four patches) for the 48 onset scenarios. Our results demonstrate that estuarine incidence in each patch tends to underestimate the entire coastline incidence, but estuarine populations in patch 2 are most representative (Figure 5C and S15). This is particularly the case for the seven high risk scenarios. Thus, we suggest that sentinel surveillance for DMV would be most effective if carried out in estuarine individuals occupying patch 2.

## 3.0 Discussion

Our data-driven estimates of both movement and contact rates in Tamanend’s bottlenose dolphins highlight differences in behavior along the coastline and inform the role of metapopulation structure on their vulnerability to infectious disease threats. We find that individual movement along the Atlantic coast differs spatially and by ecotype. We also highlight the use of our model in optimizing epidemic surveillance and risk assessments in this vulnerable population.

We found that movement rates are higher between the southern metapopulation patches compared to the northern patches, which could be partially explained by the estimates of seasonal stock presence in these regions. For example, while only one migratory coastal stock is thought to move between patches 1 and 2, there are two coastal stocks thought to move between the southern patches^25^. There was also more variation in coastal movement rates compared to estuarine which could be due to the smaller sample size of coastal individuals, or temporal differences in movement between the two ecotypes, as estuarine populations tend to engage in a more year-round dispersal while coastal populations engage in seasonal migration^25^. We also found that coastal individuals moved at a slower rate in the north which was important in reproducing infection dynamics similar to the past two DMV outbreaks; if coastal individuals migrate too quickly into the northern coastline (e.g., Delaware and New Jersey), infection originating further south (e.g., Virginia or North Carolina) wouldn’t reach this area. This suggests that factors that affect the rates of dolphin migration could have significant impacts on future epidemics. For example, recent poleward shifts of odontocete communities in the Northeast United States^50,51^ could indicate increased migration into higher latitudes which could subsequently affect the dynamics of contagious infections.

We find that increased breathing synchrony contact in the breeding season results in epidemic dynamics consistent with past outbreak data. This type of contact, which is important for respiratory disease spread in cetaceans and varies based on age and sex class^36^, is also higher in coastal individuals than estuarine individuals, which can explain a higher coastal infection burden. This difference could be due to a variety of factors. First, coastal population^25^ and group^52^ size estimates tend to be much larger than estuarine estimates; stock assessment reports put coastal population sizes in the thousands, while estuarine estimates are in the hundreds^25^. These larger sizes could provide a greater opportunity for more contact among coastal ecotypes (assuming density-dependent contact). Second, coastal individuals tend to have much larger habitat ranges^25^, which may result in a large allocation of their activity budgets to traveling, which commonly occurs in groups with greater opportunity for synchronized breathing. Further, being farther offshore in a more open ocean gives coastal individuals greater access to large schooling fish, which might require foraging to be more social compared to estuarine environments where it is often more advantageous to be solitary^53^. While NOAA has declared two coastal stocks depleted due to DMV infections^39^, no estuarine stocks have the same designation. However, our predictions show that nearly 50% of estuarine populations would be infected across the coastline. The true effect of DMV on estuarine individuals is unclear^25,47^, but uncovering these effects is especially important as many mid-Atlantic estuarine stocks are considered strategic due to the impact of fisheries interactions^25^. Enhanced surveillance for the presence of DMV antibodies in estuarine individuals^26^ could provide a clearer picture of the true impact of DMV on estuarine ecotypes.

Our model not only demonstrates the role of metapopulation structure on disease dynamics, but can also be applied to improve preparedness for future disease outbreaks. We examined epidemic risk along the coastline if outbreaks began in a different time and place than observed in the past, and found that only seven scenarios carry the same risk as historic DMV outbreaks, the majority of which begin at northern latitudes. This suggests that dolphin behavior likely structures epidemic dynamics in a way that results in emergence hotspots in the north which could explain why the 1987 and 2013 outbreaks were so similar in their dynamics. Ecologically, individuals at northern latitudes may also be at high risk for initial infections as they likely have more chances for interactions with other cetacean species that act as reservoirs for morbillivirus, such as long finned pilot whales (*Globicephala melas*)^54,55^. However, only scenarios that began in migratory season 1 (April-June) produced infection dynamics similar to historic DMV epidemics. Our model revealed that at least four scenarios where outbreaks began in migratory season 2 (October-December) carried high epidemic risk but resulted in different dynamics, suggesting a potential for future DMV epidemics that peak in the south during the cold water season. This finding is critically important to the development of both control and surveillance strategies as it points to seasons and locations where these measures might be most effective.

While the control of respiratory disease spread in dolphins remains a challenge, surveillance of DMV in Tamanend’s dolphins is feasible through boat based biopsies^26^ particularly in individuals of estuarine ecotype given their spatial ecology. Our work shows that estuarine individuals in patch 2 (i.e., individuals in the northern North Carolina estuarine stock) would be optimal sentinels for routine DMV surveillance for all Tamanend’s dolphins during or following an outbreak. Indeed, past DMV surveillance work carried out in Georgia (our patch 4) showed a low proportion of estuarine individuals infected compared to coastal individuals^26^, indicating that estuarine stocks in this region may not be the best indicators of infection during or post epidemic.

Metapopulation disease models parameterized with fine-scale host movement patterns have been theoretically illustrated to be critical in predicting geographically heterogeneous disease dynamics^56^. As our model captures spatial and ecotype heterogeneity in movement and contact among a complex marine species, we demonstrate that such empirically-informed structured models indeed reproduce observed dynamics compared to models without such variation. This suggests that such biological processes are critically important to consider for understanding disease spread. Our results also provide empirical support for theoretical metapopulation disease models, which show that while highly contagious diseases can spread widely through well connected metapopulations^57^, infection tends to decline in subsequent patches relative to the origin patch^58^. Indeed, while infection spreads to all patches in our model, the highest number of infections occur in the origin patch (2) with declining infection rates into the subsequent patches.

Our model can be expanded upon and applied to forecast future epidemics in a variety of scenarios. First, working with a collated dataset allows us to update our model parameters as researchers contribute additional sighting data in the future. For example, considering that our sample size of coastal individuals is low compared to their estimated population sizes (Table S3), obtaining new data from current research sites could allow us to improve upon our movement rate estimates, or break up patches to examine more specific areas of the coastline^59^. Additionally, while the Florida coastline was undoubtedly impacted by DMV^22^, the current lack of data on dolphin movement in this region makes it difficult to uncover the mechanisms of infection in this area. While our supplemental analysis suggests that there is likely migration of individuals into this part of the coastline, obtaining data from researchers in Florida would allow us to expand the model to empirically include this part of the coastline. Second, we can forecast the timing of future outbreaks by incorporating life history details of dolphins such as natural birth and mortality rates^60,61^. Since an epidemic is typically extinguished when the number of susceptible individuals remaining is too low for infection to persist (due to death or acquired immunity), the consistent replenishment of susceptibles born into a population makes recurrent epizootics possible^62^. The 26-year lag between the DMV outbreaks in Tamanend’s dolphins^21,22^ and similar lags in deadly morbillivirus outbreaks in other dolphin^18^ and seal^63^ populations highlights the importance of predicting these interepidemic periods. Third, we could account for infection induced behavioral changes that reduce social interactions due to avoidance^64^, or sickness behaviors^65^. For example, during the 2013 DMV outbreak, network connectedness was reduced compared to connectedness pre- and post-outbreak in Tamanend’s dolphins in Florida which likely mitigated some of the spread of DMV in this region^27^. Fourth, we can account for differences in the rate of coastal and estuarine mixing along the coastline. Although we assume that our estimated ecotype mixing rate is uniform spatiotemporally, current research suggests it likely varies. For instance, in Florida, suspected estuarine dolphins have greater overlap with suspected coastal individuals in both the warm and cold water seasons compared to migratory seasons^27^, and the level of this overlap greatly exceeds that observed near Brunswick, Georgia^26^. Finally, this model can consider the role of global change on disease consequences in Tamanend’s bottlenose dolphins. For example, climatic changes such as warming oceans can thermally stress animals, making them more susceptible to disease^66^, or affect their seasonal connectivity and distribution^50^. By modifying parameters in our model to reflect changes such as these, we can make predictions as to how these factors affect disease transmission in this, and related, species^67,68^. In addition to global change, the biodiversity crisis has elevated the importance of epidemic threats^69^. This is especially true for marine mammals who are wide-ranging and transmit deadly viruses within and across species^70,71^. Models using real-world data are vital tools in the arsenal of sustained efforts to mitigate these impacts.

## 4.0 Methods

### 4.1 Metapopulation structure inference

Given the diverse movement patterns exhibited by Tamanend’s dolphin ecotypes^25^, we use the broader term ‘movement’ to describe both estuarine dispersal and coastal migration along the coastline. We defined ‘movement rate’ as the daily proportion of dolphins transitioning from one patch to another. To establish the metapopulation structure of Tamanend’s bottlenose dolphins, we allowed dolphins to move between different *patches* of coastline between New Jersey and Georgia, and contact each other within these patches. While dolphins along the Florida coastline were also significantly impacted by DMV, in this region there is a lack of the consistent sighting data needed to be included in our empirical model. Thus while we dropped this region from our analysis, we assessed this area theoretically in the Supplementary Information, section 10.1.

We allowed movement rates between patches to vary along the coastline. We also considered the differences in movement and contact between the two main *ecotypes* of Tamanend’s dolphins: *coasta*l and *estuarine*, with coastal individuals engaging in long-distance seasonal migration and estuarine animals engaging in short-distance regular dispersal. Additional details on bottlenose dolphin stock structure and size on the US Atlantic coast can be found in the Supplementary Information, sections 2.2 and 2.3.

#### 4.1.1 Defining patches and movement rates

##### Dataset

The Mid-Atlantic Bottlenose Dolphin Catalog (MABDC)^72^ is a cooperative catalog maintained by a curator containing photographs of bottlenose dolphin dorsal fins captured at different sites along the U.S. Atlantic coast by individuals and institutions conducting independent research. Since dolphins can be individually identified through the unique shape, nicks and scars on their dorsal fins, dorsal fins photographs can be used in mark-recapture studies to track individuals over time and space^73^. The MABDC is hosted through the web-based OBIS-SEAMAP portal (https://seamap.env.duke.edu), which allows researchers to search for potential matches of dolphin dorsal fins among catalogs and generate data on spatial movement for individual dolphins across seasons.

Sighting histories for individuals in the MABDC were generated by comparing dorsal fin photographs (Figure 2A) across 28 MABDC fin catalogs located between New Jersey and Georgia. We completed 173 (out of a possible 377) of these pairwise comparisons (see Supplementary Information, section 1 for details), and compiled the associated annual sighting histories of all dolphins sighted across these catalogs (Figure 2B). As we assumed that movement of dolphins is stable across years, we disregarded the year of each sighting and only examined the season. To be included, individuals had to have 1) at least five sightings, and 2) one or more sightings in the warm water season (July-September), in the cold water season (January-March), and in either migratory season 1 (April-June) or migratory season 2 (October-December) (n = 410). Seasons were established based on 20 years of sea surface temperature data (Figure S4).

##### Defining metapopulation patches

First, we divided the coastline into ecologically relevant metapopulation patches that dolphins move between during an annual period. Since NOAA considers unique dolphin stocks to be composed of individuals with similar seasonal ranges, we performed a hierarchical clustering analysis (Ward’s method) on the spatiotemporal timeseries of our 410 dolphins to predict natural groupings according to sighting histories (Figure 2C). We identified the optimal number of clusters, *C*, to be when the size of the clusters are most uniform while their silhouette scores (mean distance between clusters) are greater than the average score of the full data set (Figure S1).

Sightings of individuals in each cluster in the warm water season, when stocks are thought to overlap the least^25^ were plotted, with the resulting ranges used to divide the coastline into *C* patches (Figure 2D). These patches were assumed to be the optimal areas from which individuals of each cluster would leave once temperatures begin to decrease, and return to when temperatures begin to increase. We validated the patch configuration by comparing them to the NOAA estimated ranges for bottlenose dolphin stocks in this region (Figure S2).

##### Determining movement rates between patches by ecotype

The comparison of our patch configuration to NOAA’s stock ranges (Figure S2) suggested that estuarine ecotypes occupy one patch year round or disperse between a maximum of two patches, whereas migratory coastal ecotypes migrate between three or more patches over the course of a year. Therefore, we classified individuals sighted in only one or two patches in the MABDC as estuarine (n =373) and individuals sighted in three or more patches as coastal (n= 37).

Next, we estimated movement rates across patches for both estuarine and coastal ecotypes. Due to considerable variability in the sampling effort across the field sites that contribute to the MABDC, traditional capture–recapture models are not realistic. To analyze these data, we fit a continuous time multistate capture-recapture model that accounts for stochasticity in detection times^59^. Specifically, we fit models to both coastal and estuarine individuals and estimated the transition rate (*r*_*s j,k*_) between patch *j* and *k*, for each ecotype *s*. For the estuarine model, we only included estuarine individuals observed in more than one patch (n = 219). Additional details of this model are in the Supplementary Information, section 4.

We calculated the movement rate between patch *j* and *k* for ecotype *s* (*η*_*s j,k*_) as:

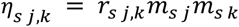

where *m*_*s j*_ and *m*_*s k*_ are the proportions of individuals of ecotype *s* in patch *j* and *k* respectively that have been seen moving to another patch as observed in the MABDC (Figure 2E).

#### 4.1.2 Defining contact rates by ecotype

##### Data Collection

To estimate the rate of disease spreading contact in bottlenose dolphins, we used behavioral data collected from boat-based observational surveys and focal animal sampling^74^ from one research site that contributes to the MABDC, the Potomac Chesapeake Dolphin Project (PCDP). The PCDP collects data in the Potomac River and middle Chesapeake Bay (patch 1) and individuals seen at this field site have been established by additional sightings in the MABDC to be both suspected estuarine and suspected coastal. Given this and the robust similarities among *Tursiops* populations, we assumed that data on contact rates obtained from this field site are representative of social contact for both ecotypes of Tamanend’s dolphins.

Since DMV is a respiratory-transmitted pathogen, we focused on synchronized breathing interactions as our measure of contact and collected the number of synchronized breaths continuously throughout a focal follow for the focal individual^73^. We collected 101 focal follows on 99 individuals between June and September 2015-2022 with an average follow time of 25 minutes.

##### Determining coastal and estuarine ecotypes for focal individuals

As none of our 101 focal PCDP individuals were sighted at another location in the MABDC, we used a different approach to classify each individual as estuarine or coastal ecotype^75^. We used a K-means clustering algorithm to cluster focal individuals based on the average distance each individual was sighted from shore which resulted in three clusters: a “nearshore” cluster (n=55 follows), a “midshore” cluster (n=30 follows), and a “farshore” cluster (n=16 follows). Based on expert opinion^25,52^, we designated individuals in our nearshore cluster as estuarine and individuals in the farshore cluster as coastal (see Supplementary Information, section 5.1 for more methodological details). Individuals in the midshore cluster were classified as “undetermined” and therefore dropped from further analysis, but a sensitivity analysis including these individuals can be found in the Supplementary Information, section 5.3.

Using our focal follow data, we estimated average daily synchrony degree for both estuarine and coastal ecotypes, which represents the number of unique individuals an animal has synchronous breathing contact with over an average day for ecotype *s, k*_*s*_, based on methodology adapted from past work^36^ (see Supplementary Information, section 5.2).

##### Calculating transmission rate *β*

We estimated the transmission rates for DMV within each ecotype *s, β*_*ss*_ (Figure 2F) based on the average synchrony degree by ecotype, *k*_*s*_ and the per contact infectiousness of DMV (*τ*= 0.032, calculated in the Supplementary Information, section 6.1):

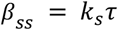

We also calculated a range for *β*_*CE*_ (Figure 2F), the transmission rate between coastal, *C*, and estuarine, *E*, ecotypes as:

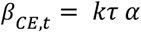

based on the average synchrony degree across both ecotypes, *k*, and the degree of mixing between estuarine and coastal stocks (*α*) estimated to vary between 0 and 6% from PCDP sighting data (see Supplementary Information, section 6.2).

### 4.2 Epidemiological metapopulation model

Using our inferred metapopulation structure, we modeled the spread of DMV in Tamanend’s dolphins between New Jersey and Georgia. Our patches were populated with individuals of both coastal and estuarine ecotypes based on the best estimates of NOAA stock assessments from before the 2013 DMV epidemic (see Supplementary Information, section 2.3). To represent migration, coastal dolphins move among patches (based on *η*_*C,jk*_) during the first (March-June) and second (October-December) migratory season and did not move in the warm (July-September) or cold (January-February) water season. To represent dispersal, estuarine dolphins move (based on *η*_*E,jk*_) a maximum of one patch south between January and June and one patch north between July and December. Within each patch we considered a standard SIR metapopulation model to simulate the spread of DMV through a patch (Figure 2G) that was discrete, stochastic^42^, and considered frequency dependent transmission for DMV^22^: susceptible dolphins (S) became infectious (*I*) and able to transmit DMV based on transmission rates {*β*}, and become removed (*R*) with removal rate *μ* based on the average infectious period of DMV (8 days)^22^. We completed 100 simulations of DMV spread using this model. Full model details can be found in the Supplementary Information, section 7.

### 4.3 Comparing model predictions to past DMV outbreaks

#### Dataset

We used stranding data consisting of records of bottlenose dolphins that washed ashore or were found floating, defined as “stranded” from March 2010-September 2014 from NMFS to capture dolphin mortality before and during the 2013-2014 DMV outbreak. We also obtained stranding data from the Smithsonian Division of Marine Mammal Collections from March 1984-September 1988 to capture dolphin mortality before and during the 1987-1988 DMV outbreak. Because not all detected cases of strandings would be due to DMV, we calculated excess mortality to estimate the number of deaths during the outbreak periods that exceed what would be expected based on the pre-outbreak periods. To do so, we removed average “background” mortality (such as from natural mortality or human interactions) for each latitude based on stranding data collected 3 years prior to each respective DMV outbreak^22,76^. We also estimated the probable date of infection for each individual based on the stranding observation date, and the stranding status (see Supplementary Information, section 8). We used these dates to estimate a time series for observed infections, hereafter referred to as “outbreak data”.

To compare our model predictions to the DMV outbreak data, we normalized the infection incidence for each patch and for each week to the maximum number of infections across all patches and all weeks. To quantify the comparison, we relied on Pearson’s correlations between both time series (higher values indicate more consistent model estimates). We additionally also measured the sum squared error (SSE), the sum of the differences between infection incidence of the model estimates and the outbreak data (lower SSE values indicate more consistent model estimates).

#### Determining the most likely start time and location of infection

NOAA had declared that DMV unusual mortality events began in July for both the 1987 and 2013 epidemics, and the first of these mortalities appeared in waters between New Jersey to Virginia. To identify likely onset locations and times for the DMV epidemic, we used Pearson’s correlations and least squares fitting (by minimizing the SSE) with the infection time series, constrained to dates between May 1st and June 30th and locations limited to patch 1 or patch 2.

#### Assessing seasonal differences in *β*

We tested two seasonal hypotheses that could further affect the spread of infection along the coastline and between coastal and estuarine stocks. (See Supplementary Information, section 6.3 for additional methodological details):

1. Seasonal social behavior: The PCDP has observed a peak in calf birth dates between April 15 and July 7 (Figure S7). Given a dolphin’s 12 month gestation period, these months are likely important for breeding in Tamanend’s dolphins. Group sizes are generally larger during the breeding season, as found in bottlenose dolphins (*Tursiops aduncus*)^77^ and other delphinid species^78,79^. Synchronized breathing rates (particularly between males) are also known to be higher during breeding seasons when males are consorting females^49^. Past work has also seen that the synchrony behavior of adult male Tamanend’s dolphins can drive epidemics^36^ suggesting that an increase in these rates could further exacerbate population level outcomes. Therefore, since breeding and synchrony rates are closely linked, we consider the effect a reduced transmission rate (*β*) outside of April 15th to July 7th to account for a lower synchrony degree.
2. Seasonal environment: Similar to other enveloped viruses (SARS-CoV-2, influenza), morbillivirus is likely to survive longer in colder air and is rapidly inactivated in warm temperatures^48^. This could indicate that DMV is more efficient at transmitting, and thus more infectious, in colder seasons. Therefore, we reduced transmission rates (*β*) outside of the cold water season of January-March to account for lower per contact DMV infectiousness (*τ*).

We performed a sensitivity analysis to assess how adjusting 1) the length of the breeding or cold water season and 2) the amount by which *β* is reduced outside of these seasons affected model results (see Supplementary Information, section 6.4).

### 4.4 Evaluating how mechanisms of metapopulation structure impact the spread of DMV

Using our model, we considered how different components of metapopulation structure impacted the spread of DMV by evaluating the results of three different models in which we controlled variation in: 1) movement, 2) ecotype movement, and 3) ecotype contact.

#### Movement control

This control model assumed one *η*_*E*_ range for estuarine individuals and one *η*_*C*_ range for all coastal individuals among all patches by taking the average of the corresponding *η* values. This model removes the differences in movement rates along the coastline but retains the differences in movement rates between coastal and estuarine ecotypes.

#### Ecotype movement control

This control model assumes the same *η* range for both coastal and estuarine individuals between all patches which removes the observed differences in movement rate both along the coastline and by ecotype.

#### Ecotype contact structure

This control model assumes *β*_*EE*_=*β*_*CC*_ by averaging the two ranges which removes the variability in contact among ecotypes.

### 4.5 Applying the model to inform preparedness for future DMV epidemics

We use our model to inform 1) how the epidemic risk of DMV varies depending on the start time and location of a DMV outbreak, and 2) the best patch for establishing surveillance sites for DMV in Tamanend’s bottlenose dolphins.

#### Epidemic risk analysis

Epidemic risk for an infection was calculated beginning in each month of the year (twelve), in each patch (four) for 48 total scenarios. For each scenario, we simulate 100 epidemics, and calculate epidemic risk as the epidemic size (the proportion of individuals that ended up infected across all patches) multiplied by the epidemic probability (the proportion of simulations with an epidemic size was greater than 10%). We also examined the SSE and Pearson’s correlation values for all scenarios to the 2013 outbreak data to determine their similarity to historic DMV outbreaks in Tamanend’s dolphins.

#### Sentinel surveillance analysis

NMFS has established management units of estuarine stocks in patches 2, 3, and 4. Since surveillance methods are more practical to carry out on estuarine stocks than coastal stocks, we examined the difference in weekly infection incidence between the estuarine individuals in these three patches (normalized by the total number of estuarine individuals in the patch), and the infection incidence along the full coastline (normalized by the total number of individuals across patches and ecotypes). The resulting difference is thus a measure of how much estuarine incidence in each patch over or underestimates total coastline incidence; the patch where this difference is closest to zero for all 48 epidemic scenarios would be the best option for establishing a sentinel surveillance site.

## Acknowledgements

We thank the following contributors of the Mid-Atlantic Bottlenose Dolphin Catalog, without whom this work would not have been possible: Annie Gorgone, Mike Gould, Tara Cox, Robin Perrtree, Brian Balmer, Keith Rittmaster, Rich Mallon-Day, Todd Speakman, Eric Zolman, Jessica Taylor, Melissa Laurino, Jacalyn Toth, Daniela da Silva, Jessica Conway, Adam Fox, Justin Greenman, and Amy Engelhaupt. We also thank Ei Fujioka from Duke’s Marine Geospatial Ecology Laboratory for building and maintaining the MABDC OBIS-Seamap application. We thank the Potomac-Chesapeake Dolphin Project field assistants for 2018-2022 without whom contact data collection would not have been possible: Marley Dooling, Meng-Chun Grace Chung, Amelia Smith, Katie Knotek, Sophie Hanson, Milan Dolezal, Trevyn Toone, Molly Albright, Haley Land-Miller, Casey Marker, Kelsey O’Donnell, Jessica Wang, Jazmin Garcia. We thank the members of the Greater Atlantic and Southeast Regional Marine Mammal Stranding Networks for the collection of the stranding data used, and the National Marine Fisheries Service Marine Mammal Health and Stranding Response Program for providing this data. All PCDP data were collected under GU Permit #s IACUC-13-069, 07-041, 10-023; 2016-1235 and under NMFS permit nos 19403 and 23782.

## References

1. McCallum, H. & Dobson, A. Detecting disease and parasite threats to endangered species and ecosystems. Trends Ecol. Evol. 10, 190–194 (1995).

2. Daszak, P., Cunningham, A. A. & Hyatt, A. D. Emerging Infectious Diseases of Wildlife--Threats to Biodiversity and Human Health. Science 287, 443–449 (2000).

3. Bossart, G. D. Marine Mammals as sentinel species for ocean and human health. Oceanography 19, 134–137 (2006).

4. Heithaus, M. R., Frid, A., Wirsing, A. J. & Worm, B. Predicting ecological consequences of marine top predator declines. Trends Ecol. Evol. 23, 202–210 (2008).

5. Moore, S. Marine mammals as ecosystem sentinels. J. Mammology 89, 534–540 (2008).

6. Sanderson, C. E. & Alexander, K. A. Unchartered waters: Climate change likely to intensify infectious disease outbreaks causing mass mortality events in marine mammals. Glob. Change Biol. 26, 4284–4301 (2020).

7. Altizer, S., Bartel, R. & Han, B. A. Animal Migration and Infectious Disease Risk. Science 331, 296–303 (2011).

8. Sah, P., Mann, J. & Bansal, S. Disease implications of animal social network structure: A synthesis across social systems. J. Anim. Ecol. 1–13 (2018) doi:10.1111/1365-2656.12786.

9. Cassirer, E. F. et al. Pneumonia in bighorn sheep: Risk and resilience. J. Wildl. Manag. 82, 32–45 (2018).

10. Blanchong, J. A. et al. Respiratory Disease, Behavior, and Survival of Mountain Goat Kids. J. Wildl. Manag. 82, 1243–1251 (2018).

11. Hochachka, W. M. & Dhondt, A. A. Density-dependent decline of host abundance resulting from a new infectious disease. Proc. Natl. Acad. Sci. 97, 5303–5306 (2000).

12. O’Dea, M. A., Jackson, B., Jackson, C., Xavier, P. & Warren, K. Discovery and Partial Genomic Characterisation of a Novel Nidovirus Associated with Respiratory Disease in Wild Shingleback Lizards (Tiliqua rugosa). PLOS ONE 11, e0165209 (2016).

13. Sumithra, T. G. et al. Mycoplasmosis in wildlife: a review. Eur. J. Wildl. Res. 59, 769–781 (2013).

14. Leung, N. H. L. Transmissibility and transmission of respiratory viruses. Nat. Rev. Microbiol. 19, 528–545 (2021).

15. Duignan, P. J. et al. Phocine Distemper Virus: Current Knowledge and Future Directions. Viruses 6, 5093–5134 (2014).

16. Van Bressem, M. F. et al. Cetacean morbillivirus: current knowledge and future directions. Viruses 6, 5145–5181 (2014).

17. Aguilar, A. & Raga, J. A. The Striped Dolphin Epizootic in the Mediterranean Sea. Ambio 22, 524–528 (1993).

18. Keck, N. et al. Resurgence of Morbillivirus infection in Mediterranean dolphins off the French coast. Vet. Rec. 166, 654–655 (2010).

19. Rubio-Guerri, C. et al. Unusual striped dolphin mass mortality episode related to cetacean morbillivirus in the Spanish Mediterranean sea. BMC Vet. Res. 9, 106 (2013).

20. Costa, A. P. B., Mcfee, W., Wilcox, L. A., Archer, F. I. & Rosel, P. E. The common bottlenose dolphin (Tursiops truncatus) ecotypes of the western North Atlantic revisited: an integrative taxonomic investigation supports the presence of distinct species. Zool. J. Linn. Soc. 196, 1608–1636 (2022).

21. Lipscomb, T. P., Schulman, F. Y., Moffett, D. & Kennedy, S. Morbilliviral disease in Atlantic bottlenose dolphins (Tursiops truncatus) from the 1987-1988 epizootic. J. Wildl. Dis. 30, 567–571 (1994).

22. Morris, S. E. et al. Partially observed epidemics in wildlife hosts: modelling an outbreak of dolphin morbillivirus in the northwestern Atlantic, June 2013–2014. J. R. Soc. Interface 12, (2015).

23. Norman, S. A. et al. The application of GIS and spatiotemporal analyses to investigations of unusual marine mammal strandings and mortality events. Mar. Mammal Sci. 28, E251–E266 (2012).

24. Dougherty, E. R., Seidel, D. P., Carlson, C. J., Spiegel, O. & Getz, W. M. Going through the motions: incorporating movement analyses into disease research. Ecol. Lett. 21, 588–604 (2018).

25. Hayes, S. A. et al. U.S. Atlantic and Gulf of Mexico Marine Mammal Stock Assessments 2022. NOAA Tech Memo NMFS NE 304 (2023).

26. Balmer, B. et al. Ranging patterns, spatial overlap, and association with dolphin morbillivirus exposure in common bottlenose dolphins (Tursiops truncatus) along the Georgia, USA coast. Ecol. Evol. 8, 12890–12904 (2018).

27. Szott, E. A., Brightwell, K. & Gibson, Q. Assessment of social mixing and spatial overlap as a pathway for disease transmission in a northeast Florida estuarine dolphin community. Mamm. Biol. 102, 1267–1283 (2022).

28. Weber, N. et al. Badger social networks correlate with tuberculosis infection. Curr. Biol. 23, R915–R916 (2013).

29. Hamede, R. K., Bashford, J., McCallum, H. & Jones, M. Contact networks in a wild Tasmanian devil (Sarcophilus harrisii) population: using social network analysis to reveal seasonal variability in social behaviour and its implications for transmission of devil facial tumour disease. Ecol. Lett. 12, 1147–1157 (2009).

30. Rendell, L., Cantor, M., Gero, S., Whitehead, H. & Mann, J. Causes and consequences of female centrality in cetacean societies. Philos. Trans. R. Soc. B Biol. Sci. 374, 20180066. (2019).

31. Wells, R. S., Scott, M. D. & Irvine, A. B. The Social Structure of Free-Ranging Bottlenose Dolphins. in Current Mammalogy (ed. Genoways, H.) vol. 1 247–305 (Plenum Press, New York and London, 1987).

32. Mann, J. Behavioral sampling methods for cetaceans: a review and critique. Mar. Mammal Sci. 15, 102–122 (1999).

33. Brager, S. Diurnal and seasonal behavior patterns of bottlenose dolphins (Tursiops truncatus). Mar. Mammal Sci. 9, 434–438 (1993).

34. Sakai, M., Morisaka, T., Kogi, K., Hishii, T. & Kohshima, S. Fine-scale analysis of synchronous breathing in wild Indo-Pacific bottlenose dolphins (Tursiops aduncus). Behav. Process. 83, 48–53 (2009).

35. Moller, L. M. & Harcourt, R. G. Social dynamics and activity patterns of bottlenose dolphins, Tursiops truncatus, in Jarvis Bay, S.E. Australia. Proc. Linneaen Soc. New South Wales 120, 181–189 (1998).

36. Collier, M. A. et al. Breathing in sync: how a social behavior structures respiratory epidemic risk in bottlenose dolphins. 2023.12.01.569646 Preprint at 10.1101/2023.12.01.569646 (2023).

37. 2013-2015 Bottlenose Dolphin Unusual Mortality Event in the Mid-Atlantic (Closed). Marine Life in Distress (2018).

38. Mclellan, W. A., Friedlaender, A. S., Mead, J. G., Potter, C. W. & Pabst, D. A. Analysing 25 years of bottlenose dolphin (Tursiops truncatus) strandings along the Atlantic coast of the USA : do historic records support the coastal migratory stock hypothesis ? J. Cetacean Res. 4, 297–304 (2002).

39. NOAA Fisheries. Depleted Designation for Western North Atlantic Coastal Migratory Stock of Bottlenose Dolphins. https://www.fisheries.noaa.gov/action/depleted-designation-western-north-atlantic-coastal-migratory-stock-bottlenose-dolphins (1993).

40. Waring, G. T., Josephson, E., Maze-Foley, K. & Rosel, P. E. US Atlantic and Gulf of Mexico Marine Mammal Stock Assessments 2015. NOAA Tech Memo NMFS NE 238 501 (2016).

41. White, L. A., Forester, J. D. & Craft, M. E. Dynamic, spatial models of parasite transmission in wildlife: Their structure, applications and remaining challenges. J. Anim. Ecol. 87, 559– 580 (2018).

42. Colombi, D. et al. Mechanisms for lyssavirus persistence in non-synanthropic bats in Europe: insights from a modeling study. Sci. Rep. 9, (2019).

43. Langwig, K. E. et al. Mobility and infectiousness in the spatial spread of an emerging fungal pathogen. J. Anim. Ecol. 90, 1134–1141 (2021).

44. Fulford, G. R. & Roberts, M. G. The Metapopulation Dynamics of an Infectious Disease : Tuberculosis in Possums. Theor. Popul. Biol. 29, 15–29 (2002).

45. Brandell, E. E., Dobson, A. P., Hudson, P. J., Cross, P. C. & Smith, D. W. A metapopulation model of social group dynamics and disease applied to Yellowstone wolves. Proc. Natl. Acad. Sci. U. S. A. 118, e2020023118 (2021).

46. Swinton, J., Harwood, J., Grenfell, B. T. & Gilligan, C. A. Persistence thresholds for phocine distemper virus infection in harbour seal Phoca vitulina metapopulations. J. Anim. Ecol. 67, 54–68 (1998).

47. Scott, G. P., Burn, D. M. & Hansen, L. J. The dolphin dieoff: long-term effects and recovery of the population. in OCEANS ‘88. ‘A Partnership of Marine Interests’. Proceedings 819– 823 (IEEE, Baltimore, MD, USA, 1988). doi:10.1109/OCEANS.1988.794905.

48. Rijks, J. M., Osterhaus, A. DME., Kuiken, T. & Frolich, K. Morbillivirus Infections. in Infectious Diseases of Wild Mammals and Birds in Europe (eds. Gaivier-Widen, D., Meredith, A. & Duff, J.P.) 99–118 (John Wiley & Sons Incorporated, 2012).

49. Connor, R. C., Smolker, R. & Bejder, L. Synchrony, social behaviour and alliance affiliation in Indian Ocean bottlenose dolphins, Tursiops aduncus. Anim. Behav. 72, 1371–1378 (2006).

50. van Weelden, C., Towers, J. R. & Bosker, T. Impacts of climate change on cetacean distribution, habitat and migration. Clim. Change Ecol. 1, (2021).

51. Thorne, L. H., Heywood, E. I. & Hirtle, N. O. Rapid restructuring of the odontocete community in an ocean warming hotspot. Glob. Change Biol. 28, 6524–6540 (2022).

52. Read, A. J. et al. Stock Discrimination of Bottlenose Dolphins along the Outer Banks of North Carolina; Implications for the Risk of Entanglement in Coastal Gill Net Fisheries. Final Report North Carolina Sea Grant Bycatch Reduction Marine Mammal Project 10 DMM 01 (2013).

53. Roberts, S. M. et al. Tight spatial coupling of a marine predator with soniferous fishes: Using joint modelling to aid in ecosystem approaches to management. Divers. Distrib. 29, 1074–1089 (2023).

54. van Bressem, M.-F., Jepson, P. & Barrett, T. Further Insight on the Epidemiology of Cetacean Morbillivirus in the Northeastern Atlantic. Mar. Mammal Sci. 14, 605–613 (1998).

55. Duignan, P. J. et al. Morbillivirus infection in two species of pilot whale (globicephala sp.) from the western Atlantic. Mar. Mammal Sci. 11, 150–162 (1995).

56. Daversa, D. R., Fenton, A., Dell, A. I., Garner, T. W. J. & Manica, A. Infections on the move: how transient phases of host movement influence disease spread. Proc. R. Soc. B Biol. Sci. 284, 20171807 (2017).

57. Hess, G. Disease in Metapopulation Models: Implications for Conservation. Ecology 77, 1617–1632 (1996).

58. Michalska-Smith, M., VanderWaal, K. & Craft, M. E. Asymmetric host movement reshapes local disease dynamics in metapopulations. Sci. Rep. 12, 9365 (2022).

59. Rushing, C. S. An ecologist’s introduction to continuous-time multi-state models for capture–recapture data. J. Anim. Ecol. 92, 936–944 (2023).

60. Ben-Horin, T. et al. Modelling marine diseases. in Marine Disease Ecology (eds. Behringer, D. C., Silliman, B.R. & Lafferty, K.D.) 233–256 (Oxford University Press, 2020). doi:10.1093/oso/9780198821632.003.0012.

61. Keeling, M. J. & Rohani, P. Modeling Infectious Disease. (Princeton University Press, Princeton, 2008).

62. Grenfell, B. & Keeling, M. Dynamics of infectious disease. in Theoretical Ecology: Principles and Applications (eds. May, R. & McLean, A.) 132–147 (Oxford Press, 2007).

63. Härkönen, T. et al. The 1988 and 2002 phocine distemper virus epidemics in European harbour seals. Dis. Aquat. Organ. 68, 115–130 (2006).

64. Stockmaier, S., Ulrich, Y., Albery, G. F., Cremer, S. & Lopes, P. C. Behavioural defences against parasites across host social structures. Funct. Ecol. 37, 809–820 (2023).

65. Lopes, P. C., Block, P. & König, B. Infection-induced behavioural changes reduce connectivity and the potential for disease spread in wild mice contact networks. Sci. Rep. 6, 1–10 (2016).

66. Learmonth, J. A. et al. Potential effects of climate change on marine mammals. in Oceanography and Marine Biology: An Annual Review (eds. Gibson, R. N., Atkinson, J.A. & Gordon, J.D.M.) 431–464 (Taylor & Francis, 2006).

67. Dickinson, E. R. et al. Host movement dominates the predicted effects of climate change on parasite transmission between wild and domestic mountain ungulates. R. Soc. Open Sci. 11, 230469 (2024).

68. Gulland, F. M. D. et al. A review of climate change effects on marine mammals in United States waters: Past predictions, observed impacts, current research and conservation imperatives. Clim. Change Ecol. 3, 100054 (2022).

69. Kim, K., Dobson, A. P., Gulland, F. M. D. & Harvell, C. D. Diseases and the conservation of marine biodiversity. in Marine conservation biology (eds. Norse, E. A. & Crowder, L. B. 149–163 (Island Press, 2005).

70. Gulland, F. M. D. & Hall, A. J. Is marine mammal health deteriorating? Trends in the global reporting of marine mammal disease. EcoHealth 4, 135–150 (2007).

71. Simeone, C. A., Gulland, F. M. D., Norris, T. & Rowles, T. K. A systematic review of changes in marine mammal health in North America, 1972-2012: The need for a novel integrated approach. PLoS ONE 10, e0142105 (2015).

72. Urian, K., Hohn, A. A. & Hansen, L. J. Status of the Photo-Identification Catalog of Coastal Bottlenose Dolphins of the Western North Atlantic: Report of a Workshop of Catalog Contributors. (1999).

73. Hammond, P., Mizroch, S. & Donovan, G. Individual Recognition of Cetaceans: Use of Photo-Identification and Other Techniques to Estimate Population Parameters. In: Report of the International Whaling Comission). https://www.semanticscholar.org/paper/Individual-recognition-of-cetaceans%3A-use-of-and-to-Hammond-Mizroch/691e0e729d5e8d2665b34af427e38687a1ad4c2a (1990).

74. Karniski, C. et al. A comparison of survey and focal follow methods for estimating individual activity budgets of cetaceans. Mar. Mammal Sci. 31, 839–852 (2015).

75. Toth, J. L., Hohn, A. A., Able, K. W. & Gorgone, A. M. Defining bottlenose dolphin (Tursiops truncatus) stocks based on environmental, physical, and behavioral characteristics. Mar. Mammal Sci. 28, 461–478 (2012).

76. Karlinsky, A. & Kobak, D. Tracking excess mortality across countries during the COVID-19 pandemic with the World Mortality Dataset. eLife 10, e69336 (2021).

77. Mann, J., Connor, R. C., Barre, L. M. & Heithaus, M. R. Female reproductive success in bottlenose dolphins (Tursiops sp.): Life history, habitat, provisioning, and group-size effects. Behav. Ecol. 11, 210–219 (2000).

78. Daura-Jorge, F. G., Wedekin, L. L., Piacentini, V. de Q. & Simões-Lopes, P. C. Seasonal and daily patterns of group size, cohesion and activity of the estuarine dolphin, Sotalia guianensis (P.J. van Bénéden) (Cetacea, Delphinidae), in southern Brazil. Rev. Bras. Zool. 22, 1014–1021 (2005).

79. Karczmarski, L., Cockcroft, V. G. & McLachlan, A. Group size and seasonal pattern of occurrence of humpback dolphins Sousa chinensis in Algoa Bay, South Africa. South Afr. J. Mar. Sci. 21, 89–97 (1999).

