## Supplementary Information for "Seasonal contact and migration structure mass epidemics and inform outbreak preparedness in bottlenose dolphins"

#### Section 1: Methodology for dorsal fin matching among the Mid-Atlantic Bottlenose Dolphin Catalog

We used the AI software finfindR[1] which can automatically trace dorsal fins in photographs in one catalog, hereafter referred to as the "query catalog", and compare these to dorsal fin traces in another catalog, hereafter referred to as the "comparison catalog". For each fin in a query catalog, the program will rank the fins in the comparison catalog by a distance measure, where fins with lower distance measures are a more likely match to the fin in the query catalog. In the interest of time, for each fin in the query catalog, we checked the 5 most likely matches returned by finfindR from the comparison catalog, and only if they had a distance measure of less than 360 (our analysis of our first two catalog comparisons in which we checked the first 10 potential matches regardless of the distance measure showed that ~90% of matches had distance measures lower than 360). Any fin determined to be of low distinctiveness in the query catalog (e.g., clean fins with no distinctive markings) was not examined[2] as these fins are not possible to match. Any fin that was classified as "mutilated" in the query catalog, (i.e., severally chopped or unusually shaped) was compared manually to the mutilated fins in the comparison catalog (i.e., without using findfindR) as we observed poor performance of finfindR on these fin shapes. This process was carried out by a research assistant with 4 years of dolphin photo-ID experience. Any matches between catalogs made by the assistant were then checked and submitted to the MABDC by a researcher with 7 years of dolphin photo-ID experience. Finally, each match was then confirmed by the lead contributors for both the query and comparison catalogs (Table S2), and the MABDC curator to be included in this analysis.

Prior to the start of this project a total of 99 pairwise comparisons (out of a possible 377) had already been completed by the MABDC curator and contributors. These comparisons did not follow the same protocol, but instead were done manually or if using finfindR, looked at a greater number of potential matches for each fin. Since our methods were less in-depth than those of other researchers, we assume that by not redoing their comparisons with our methodology we will not miss any matches. However, while this could indicate that our methods may result in finding less matches than other protocols, previous analysis[1] suggests that our methodology would obtain ~90% of all the matches that exist in a comparison indicating that this loss should not be significant.

We ranked the remaining 278 comparisons by order of priority based on the proximity of the field sites where those that are farther from each other (e.g., New Jersey and Georgia) were a lower priority for comparison. Comparisons were ranked as either high (176 comparisons), medium (110), or low (91) priority and were completed in that order. By the end of our analysis, 118 (67%) high priority, 44 (40%) medium priority and 11 (12%) low priority comparisons were completed (Table S1). Based on NOAA stock structure, the chance of finding matches in the medium and low priority comparisons is low, so we assume that not completing these comparisons will have no effect on our results.

**Table S1: The status of dorsal fin comparisons among 28 catalogs in the MABDC**. Each row and column represents a catalog (details found in Table S2). Black X's indicate that the comparison was completed before the start of this project. Blue X's indicate the comparison was completed through methodology explained in this work. Green boxes indicate high priority comparisons, orange medium priority, and red low priority.

| | NJ1 | NJ2 | NJ3 | NJ4 | NJ5 | NJ6 | DE | VA1 | VA2 | MD | NC1 | NC2 | NC3 | NC4 | NC5 | SC1 | SC2 | SC3 | SC4 | SC5 | SC6 | SC7 | SC8 | SC9 | GA1 | GA2 | GA3 | GA4 |
|---|---|---|---|---|---|---|---|---|---|---|---|---|---|---|---|---|---|---|---|---|---|---|---|---|---|---|---|---|
| NJ1 | | X | X | X | X | X | | X | X | | X | X | X | X | X | X | | | | | | | X | X | X | X | X | |
| NJ2 | | | X | X | X | X | | X | X | | X | X | X | X | X | | | | | | | | X | X | X | X | X | |
| NJ3 | | | | X | X | X | X | X | X | | | X | | X | | | | | | | | | | | | | | |
| NJ4 | | | | | X | X | | X | X | X | X | X | X | X | | | | | | | | | | | | | | |
| NJ5 | | | | | | X | X | X | X | | X | X | X | X | X | | | | | | | | | | | | | |
| NJ6 | | | | | | | X | X | X | | X | X | X | X | X | | | | | | | | | | | | | |
| DE | | | | | | | | X | X | X | | | | | | | | | | | | | | | | | | |
| VA1 | | | | | | | | | | X | X | X | | | | | | | | | | | | | | | | |
| VA2 | | | | | | | | | | X | X | X | X | X | X | | | | | | | | | | | | | |
| MD | | | | | | | | | | | X | X | | X | | | | | | | | | | | | | | |
| NC1 | | | | | | | | | | | | X | X | X | X | | | | | | | | | | | | | |
| NC2 | | | | | | | | | | | | | X | X | X | X | | | | | | | | | | | | X |
| NC3 | | | | | | | | | | | | | | X | X | | | | | | | | | | | | | X |
| NC4 | | | | | | | | | | | | | | | X | X | | | X | X | X | X | X | X | X | X | X | X |
| NC5 | | | | | | | | | | | | | | | | X | | | | | | | | | | | | |
| SC1 | | | | | | | | | | | | | | | | | X | | X | X | X | X | X | X | X | X | X | X |
| SC2 | | | | | | | | | | | | | | | | | | X | X | X | X | X | X | X | X | X | X | X |
| SC3 | | | | | | | | | | | | | | | | | | | X | X | X | X | X | X | X | X | X | X |
| SC4 | | | | | | | | | | | | | | | | | | | | | | X | X | X | X | X | X | X |
| SC5 | | | | | | | | | | | | | | | | | | | | | X | | | | | | | |
| SC6 | | | | | | | | | | | | | | | | | | | | | | X | X | X | X | X | X | X |
| SC7 | | | | | | | | | | | | | | | | | | | | | | | X | X | X | X | X | X |
| SC8 | | | | | | | | | | | | | | | | | | | | | | | | X | X | X | X | X |
| SC9 | | | | | | | | | | | | | | | | | | | | | | | | | X | X | X | X |
| GA1 | | | | | | | | | | | | | | | | | | | | | | | | | | X | X | X |
| GA2 | | | | | | | | | | | | | | | | | | | | | | | | | | | X | X |
| GA3 | | | | | | | | | | | | | | | | | | | | | | | | | | | | X |
| GA4 | | | | | | | | | | | | | | | | | | | | | | | | | | | | |

**Table S2. Full names and lead contributors for all 28 MABDC catalogs included in this analysis.**

| Catalog Abbreviation from Table S1 | Full MABDC Catalog Name | Lead Contributor(s) |
|---|---|---|
| NJ1 | NJ-JLT | Jacalyn Toth |
| NJ2 | NJ-SU | Jacalyn Toth |
| NJ3 | NJ-CMWWRC | Melissa Laurino |
| NJ4 | NJ-RMD | Rich Mallon-Day |
| NJ5 | NJ-KR | Keith Rittmaster |
| NJ6 | NJ-NOAA | Annie Gorgone |
| DE | DE-CMWWRC | Melissa Laurino |
| MD | MD-PCDP | Ann-Marie Jacoby; Janet Mann |
| VA1 | VA-NOAA | Annie Gorgone |
| VA2 | VA-HDR | Amy Engelhaupt |
| NC1 | NC-RMD | Rich Mallon-Day |
| NC2 | NC-OBXCDR | Jessica Taylor |
| NC3 | NC-NCMM | Keith Rittmaster |
| NC4 | NC-DUML-UNCW | Kim Urian |
| NC5 | NC-NOAA | Annie Gorgone |
| SC1 | SC-CCU | Daniela da Silva |
| SC2 | SC-CCU-CR-JS | Daniela da Silva |
| SC3 | SC-CCU-NSC | Daniela da Silva |
| SC4 | SC-CCU-HH-AF | Daniela da Silva; Adam Fox |
| SC5 | SC-CCU-HH-JC | Daniela da Silva; Jessica Conway |
| SC6 | SC-CCU-ACE | Daniela da Silva |
| SC7 | SC-Greenman | Justin Greenman |
| SC8 | SC-NOS | Todd Speakman |
| SC9 | SC-NOAA | Annie Gorgone |
| GA1 | GA-NOAA | Annie Gorgone |
| GA2 | GA-SSU | Tara Cox; Robin Perrtree |
| GA3 | GA-UNCW | Brian Balmer |
| GA4 | GA-SAILOR | Mike Gould |

#### Section 2: Additional details on patch determination, validation, and Tamanend's bottlenose dolphin stock structure

2. 1 Determining the number of metapopulation patches from hierarchical cluster results

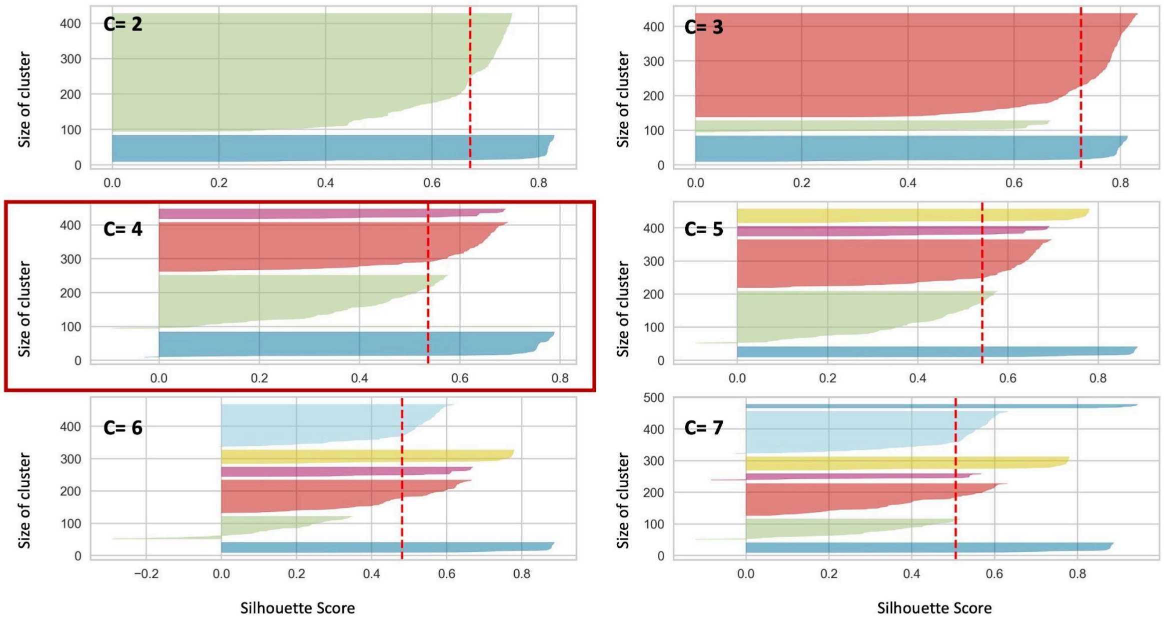


**Figure S1. Deciding the optimal number of clusters for metapopulation patch designation.** We clustered individuals based on the time series of their annual sighting histories determined with the MABDC and examined the resulting cluster sizes and silhouette scores for 2-7 clusters. We see only at two, four, and five clusters, does the silhouette score of each cluster exceed that of the average (red dashed line). Of these three, the least variability in cluster size occurs at four clusters (red box), so we choose to divide the coastline into 4 metapopulation patches.

2.2 Details on bottlenose dolphin stock structure

NOAA defines 10 stocks of two different ecotypes (three coastal and seven estuarine) of Tamanend's dolphins along the coast of the United States between New York and Georgia (Table S3) that are thought to be most delineated from each other in the warm water months[3]. As temperatures begin to drop individuals tend to disperse (estuarine) or migrate (coastal) towards the southernmost points of their habitat ranges for the cold water season where the largest amount of stock overlap is thought to occur[3] (Figure S2). Following this season, individuals will then move back to the northern parts of their habitats.

2.3 Populating patches

NOAA stock assessment reports estimate that migratory coastal stock sizes ranged between 9,000-12,000 individuals, non migratory coastal stock sizes between 4,000- 5,000, and estuarine stock sizes range between 100-1000 prior to the 2013 DMV outbreak (Table S3). We thus populate our four patches in the warm water seasons (Table S4) with individuals based on the warm water habitat ranges for the migratory coastal stocks, and the full habitat range of the non migratory and estuarine stocks (Figure S2). Since there are not consistent estimates for estuarine stocks in each patch, we assume that there are 1000 estuarine individuals in each patch.

**Table S3. The 10 stocks of dolphins recognized by NOAA between New York and Georgia and the best estimates for their size pre and post 2013 DMV outbreak.**

| Stock Name | Stock Abbreviation | Best Estimate Stock Size (Pre DMV)[4] | Best Estimate Stock Size (Post DMV)[5] |
|---|---|---|---|
| Northern Migratory Coastal | NMC | 11,548 | 6,639 |
| Southern Migratory Coastal | SMC | 9,173 | 3,751 |
| South Carolina Georgia Coastal | SCGC | 4,377 | 6,027 |
| Northern North Carolina Estuarine System | NNCES | 950 | 782 |
| Southern North Carolina Estuarine System | SNCES | 188 | UNKNOWN |
| Northern South Carolina Estuarine System | NSCES | UNKNOWN | 453 |
| Charleston Estuarine System | CES | 289 | UNKNOWN |
| Northern Georgia/Southern South Carolina Estuarine System | NGSSCES | UNKNOWN | UNKNOWN |
| Central Georgia Estuarine System | CGES | UNKNOWN | 192 |
| Southern Georgia Estuarine System | SGES | 194 | UNKNOWN |

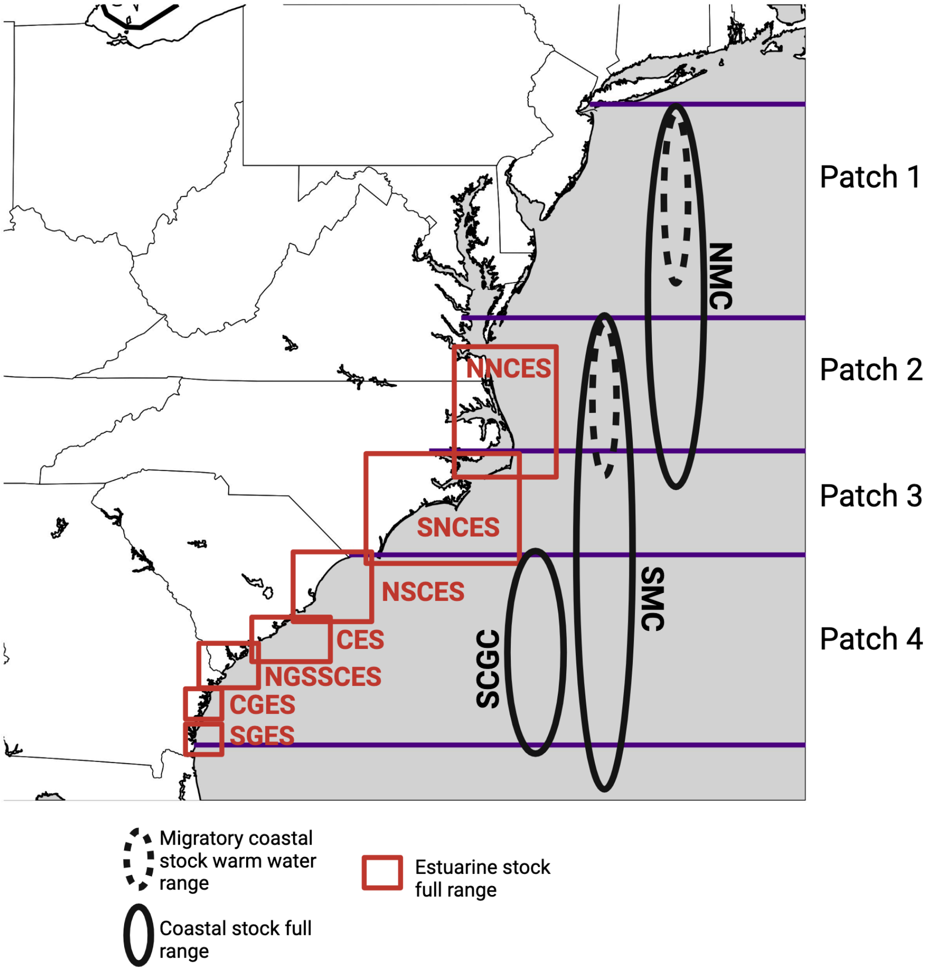


**Figure S2. Metapopulation patches with estimated habitat ranges of NOAA established stocks[3,6].** The full habitat ranges of each of the 10 stocks of Atlantic bottlenose dolphins are represented by red boxes for estuarine stocks and black circles for coastal stocks. The dashed circles show the warm water season range for the two migratory coastal stocks, demonstrating the delineation of stocks seen in this time period. We show that in the warm water season, stocks also tend to delineate into our estimated patches, providing support for our patch designation. Created with Biorender.com

**Table S4: The number of individuals in each patch in our model in the warm water season, based on NOAA stock assessment reports.**

| Patch | Number of Estuarine Individuals in Warm Season | Number Coastal Individuals in Warm Season |
|---|---|---|
| 1 | 1,000 | 10,000 |
| 2 | 1,000 | 10,000 |
| 3 | 1,000 | 2,000 |
| 4 | 1,000 | 5,000 |

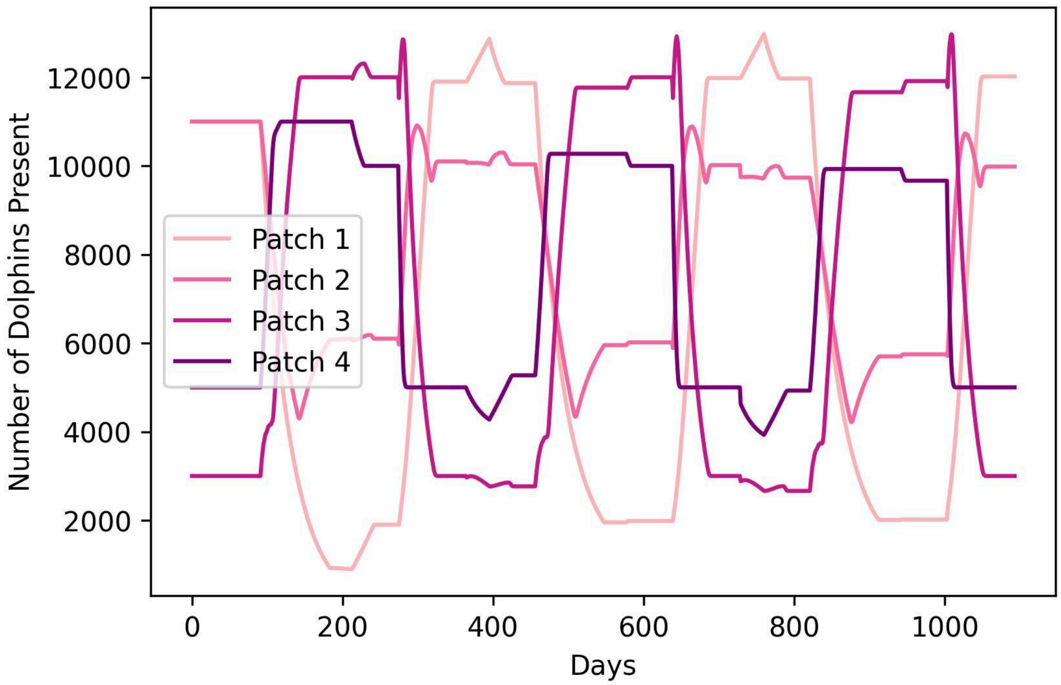


**Figure S3. Our model estimate of total patch size over the course of 18 months where day 0 is the start of the warm water season (July 1).**

#### Section 3: Determining the warm water, cold water, and migratory seasons

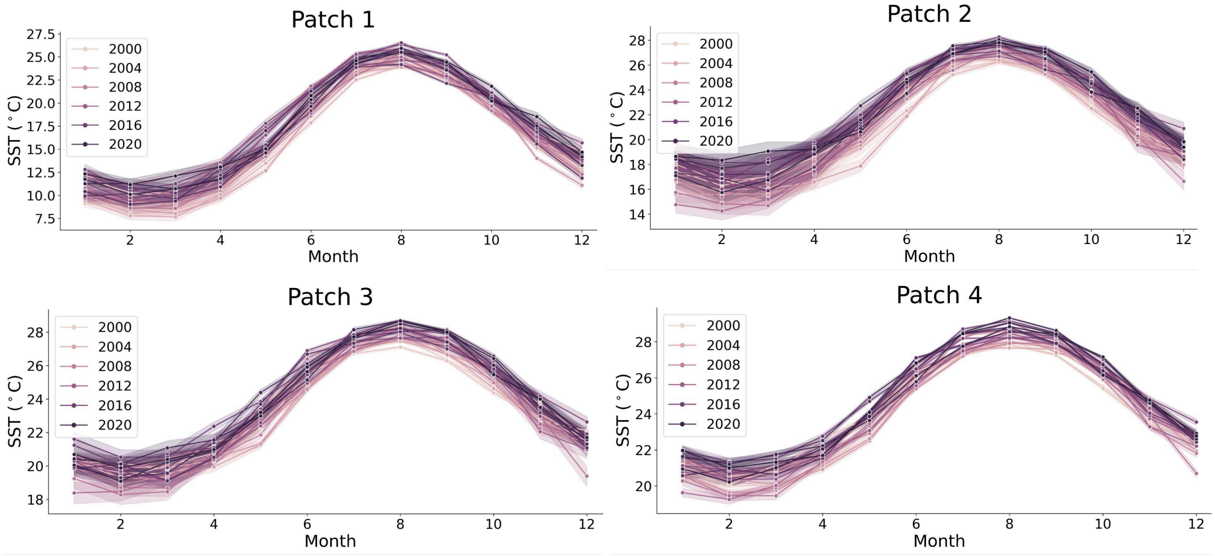


**Figure S4: The average sea surface temperature (SST) values for the coastal waters of each patch between 2000-2020.** Data obtained from the Copernicus Climate Change Service[7]. We find July, August and September to have the highest SST values across patches and years and thus determine these months to be the warm water season for our model. January, February and March have the coldest SST across patches and years and thus determine these months to be the cold water season for our model. We determine the remaining month between these seasons to be migratory season 1 (April-June) and migratory season 2 (October-December).

#### Section 4: Additional details for the continuous time multi-state capture-recapture model

The methodology as written below is adapted from[8].

First, we assumed that $i = 1, 2, ...410$ dolphins were initially photographed, and subsequently tracked using photo-ID methods that allow individuals to be detected in continuous time. Thus, each individual had a unique detection history $y_i$ of length $U_i$ indicating the time $t_{i,u}$ and patch $s_{i,u}$ of each detection $u$ where $s \in 1 - 4$ (patch identity). Detections could occur at any time between the first observation ($t_{i,0} = 0$) and the end of the study period (one year, $T = 365$). Because the individual detection histories represent a partially observed Markovian state transition process, this model should be viewed as a special case of continuous-time hidden Markov models[9,10].

Transitions between patches were governed by transition rate (4 X 4) matrix Q:

$$\begin{bmatrix} q_{11} & r_{12} & 0 & 0 \\ r_{12} & q_{22} & r_{23} & 0 \\ 0 & r_{23} & q_{33} & r_{34} \\ 0 & 0 & r_{34} & q_{44} \end{bmatrix}$$

Which describes how individuals transition from patch to patch. Off-diagonal elements ($r_{j,k}$, $j \neq k$) governed the transition rates between patches and the diagonal elements $q_{s,s}$ determined the rate at which individuals remain in their current patch. The rows of the transition rate matrix must sum to zero to function as a Markov process[11], which was ensured by setting the diagonal elements $q_{ss} = -[\sum_{j \neq k} r_{jk}]$. Since individuals can only transition between geographically connected patches, patches that are not connected had a $r_{j,k}$ of 0. The probability of transitioning between patches over the observed intervals between detections could thus be calculated as $exp\{Q\Delta t\}$ where $\Delta t$ is the length of time between consecutive detections.

Patch-specific detection rates were represented by a vector containing the detection intensities for each patch, defined as a matrix $\Lambda$ consisting of $l_i$ on the diagonal and zero otherwise:

$$\begin{bmatrix} l_1 & 0 & 0 & 0 \\ 0 & l_2 & 0 & 0 \\ 0 & 0 & l_3 & 0 \\ 0 & 0 & 0 & l_4 \end{bmatrix}$$

The patch-specific probability of detecting an individual at any length of time $\Delta t$ was thus $exp\{-\ \Lambda\Delta t\}\Lambda$.

Defining $\Delta t_{i,u}$ as $t_{i,u} - t_{i,u-1}$, $u \in [1,\ U_i]$, $Q$ and $\Lambda$ defined the probability of transitioning between states and being detected for each observation $u$. For example, if a dolphin $i$ was detected three times ($U_i$=3) in a patch, we can calculate the likelihood of each detection as:

$$\Gamma_{i,1} = exp\left\{(Q\ -\ \Lambda)\Delta t_{i,1}\right\}\Lambda$$

$$\Gamma_{i,2} = exp\left\{(Q\ -\ \Lambda)\Delta t_{i,2}\right\}\Lambda$$

$$\Gamma_{i,3} = exp\left\{(Q\ -\ \Lambda)\Delta t_{i,3}\right\}\Lambda$$

$$\Gamma_{i,4} = exp\left\{(Q\ -\ \Lambda)\Delta t_{i,4}\right\}\Lambda$$

where $\Delta t_{i,1} = t_{i,1} - t_{i,0}$, $\Delta t_{i,2} = t_{i,2} - t_{i,1}$, $\Delta t_{i,3} = t_{i,3} - t_{i,3}$, and $\Delta t_{i,4} = T - t_{i,3}$. Dolphin $i$'s individual contribution to the likelihood is thus:

$$L_i = f_i\left[\prod_{u=1}^{4} \Omega_{i,u}\right]1$$

Where $\Omega_{i,u}$ is another 4 x 4 matrix that contains only the transition and detection probabilities associated with the observed patch transitions, $f_i$ is the probability of an individual starting in each patch, and $1\ =\ 1\ 1\ 1\ 1$ .

To estimate $r_{jk}$ and $l_j$ for all patches for both estuarine and coastal individuals, we fit the above model to our n = 219 estuarine sighting histories, and again to our n=36 coastal sighting histories using the R package rstan, with code adapted from[8]. We fit four chains for 500 iterations, with a warm up period of 200 iterations, assumed initial values of all $r_{j,k}$ and $l_j$ to be 0.01 and used relatively weak gamma priors on both $r$ and $l$.

#### Section 5: Determining average daily degree for coastal and estuarine ecotypes using PCDP focal data

We collected data in Maryland and Virginia waters from the lower tidal Potomac River (southeast of the Governor Harry W. Nice Memorial Bridge, N 38.361595, W 76.997144) to the middle Chesapeake Bay (specifically Ingram Bay N 37.790233, W 76.997144) with the Potomac Chesapeake Dolphin Project (PCDP). We conducted focal follows on an individual dolphin for a predetermined amount of time between 15 minutes and 2 hours. Since DMV is a respiratory-transmitted pathogen, to estimate disease outcomes among bottlenose dolphins, we estimate synchrony degree: the number of unique individuals a focal animal has synchronous breathing interactions with over a typical day.

5.1 Assigning focal individuals to an ecotype

We calculated the average distance from shore for each individual using the coordinates of each of their established PCDP sightings between 2015-2022. We measured the distance between each sighting coordinate and a high resolution shape file of the chesapeake bay[12] that was converted to a line in R, using the st_distance function in the R package `sf`, and selecting the minimum distance. We then used a cluster analysis[13] on these average distances for all focal individuals to predict natural groupings according to their average distances from the shoreline. We used a K-means clustering algorithm ranging from one to ten clusters initialized using the k-means++ method. The within-cluster sum of squares (WCSS) was calculated for each clustering solution, and the optimal number of clusters was selected by analyzing the resulting WCSS using the elbow method. The elbow point, occurred at three clusters: a "nearshore" cluster (n=55 follows) with average distances between 37m-827 m from shore, a "midshore" cluster (n=30 follows) with average distances between 937-1864 m from shore, and a "farshore" cluster (n=16 follows) with average distances between 2004-4614 m from shore. Reports indicate that estuarine ecotypes are typically sighted less than 1 km[3] (less than 500 m in some areas[14]) from shore. Therefore, we designated individuals in our nearshore cluster as estuarine and individuals in the farshore cluster as coastal. Individuals in the midshore cluster were classified as "undetermined" and therefore dropped from further analysis, but we perform a sensitivity analysis including these individuals in section 5.3.

5.2 Calculating average daily degree for each ecotype

Since our focal follow data only measures synchronized breathing for short periods of time (average 25 minutes), it is not representative of total contact (degree) over a day. To measure relevant synchrony degree, we must therefore extrapolate our measurements for longer time periods using a previously established power law relationship between *Tursiops* synchrony degree and the number of synchronous breathes (syncs) individuals partake in[15]:

$$k_d(D) = 0.587 n_d(D)^{0.5209}$$

By estimating the average number of syncs an individual from ecotype $s$ partakes in over the course of one day $D$ =1, $n_s(D)$, we can then calculate their daily average synchrony degree $k_s(D)$.

To empirically estimate $n_s(D)$ we decompose the social processes. We define $n_s(D)$ to be a product of 1) the probability that individuals of ecotype $s$ breathe synchronously while in a group, $P_s^{sync}$, 2) the typical rate of synchronized breathing for ecotype $s$, $r_s$, and 3) the time period of interest, $D$, taken here to be one day.

$$n_s(D) = DP_s^{sync} r_s$$

Using our focal follow data, we estimate distributions for $P_s^{sync}$, and $r_s$. For $P_s^{sync}$ we use a generalized linear mixed model with a binary response variable for whether a sync occurred during a follow. We use a categorical predictor variable for ecotype, a continuous effect for the length of the follow to control for increased likelihood of syncing over time, and random effects for the year the follow was conducted and the ID of the dolphin to control for variability in data collected across years, and individual behavior of the dolphin respectively. We then choose 1000 random model estimates for each population/demographic grouping to generate a distribution for each $P_g^{sync}$. For $r_s$ we use a generalized linear mixed model with a continuous response variable for the number of syncs that occurred during a follow. We use a categorical predictor variable for ecotype and a continuous predictor for the length of the follow, and random effects for the year the follow was conducted and the ID of the dolphin. As the model estimates the number of syncs that will occur for each ecotype over the average follow length in our dataset (24.6 minutes) we calculate $r_s$ by choosing 1000 random model estimates for each ecotype and divided it by the average follow length to generate a distribution of each $r_s$.

*5.2.1 Resulting distributions:*

$$P_{coastal}^{sync} = 0.726\ [0.721 - 0.732]$$

$$P_{estuarine}^{sync} = 0.672\ [0.666 - 0.679]$$

$$r_{coastal} = 0.208\ syncs/minute\ [0.146 - 0.295]$$

$$r_{estuarine} = 0.256\ syncs/minute\ [0.210 - 0.312]$$

We estimate 1000 $n_s(D)$ for each ecotype from these distributions and calculate the average associated $k_s(D)$ from the power law fit resulting in average daily coastal degree, $k_C = 8$ and average daily estuarine degree, $k_E = 6$.

5.3 Sensitivity Analysis:

Since we dropped 30 focal follows due to undetermined stock ID, thus limiting our sample size, we conducted a sensitivity on our degree calculations using all 101 focal follows. We gave the current estimate of estuarine individuals being within 1 km of shore[3,14] a 300m buffer, so for the 30 individuals that fell into the "Undetermined" cluster, we assigned them as estuarine if their average distance from shore was less than 1.3km (new estuarine sample size n = 72), and coastal otherwise (new coastal sample size n = 29). We repeat the above analysis with these new sample sizes.

*5.3.1 Resulting distributions:*

$$P^{sync}_{coastal} = 0.822\,[0.504 - 0.954]$$

$$P^{sync}_{estuarine} = 0.756\,[0.491 - 0.909]$$

$$r_{coastal} = 0.189\; syncs/minute\,[0.143 - 0.249]$$

$$r_{estuarine} = 0.238\; syncs/minute\,[0.205 - 0.285]$$

We estimate the mean degree for both coastal and estuarine individuals based on these distributions to be 9, indicating no difference in degree between ecotypes, in contrast to when we use only our high confidence focal follows. When these average degrees are used in our disease model, estuarine and coastal infection burdens are not different and are no longer significantly biased towards coastal infections (Figure S5). While this is in contrast to expert opinion[16,17], we note that this model still returns a similar patch level infection time series (Figure S5, Table S5) to our original model (where coastal degree is higher than estuarine degree) suggesting that our model's patch level results are largely robust to potential differences in ecotype contact rates.

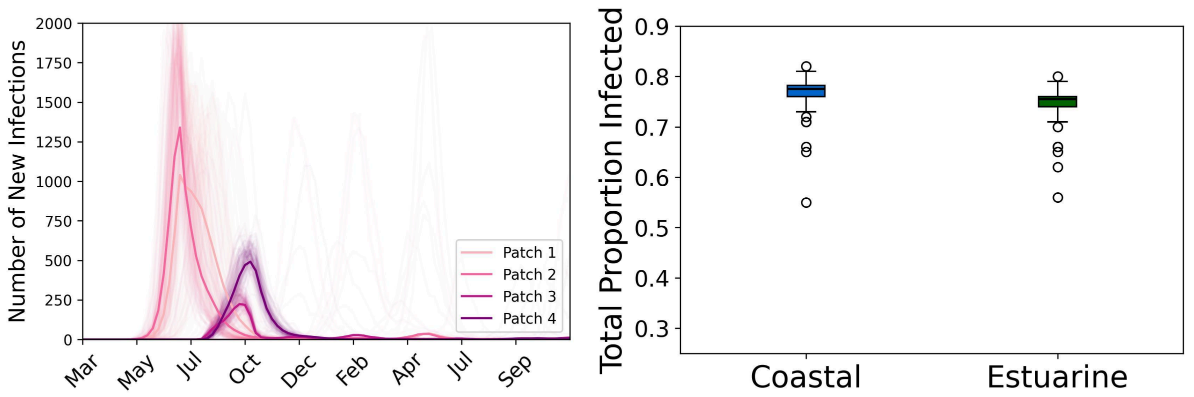


**Figure S5: The time series of new infections by patch (left) and total ecotype burden (right) when $k_C$ and $k_E$ = 9.**

**Table S5: The sum squared error (SSE) and Pearson's correlation coefficient (r) for our epidemiological disease models (based on different ecotype average degrees) compared to the 2013 outbreak data.**

| Ecotype Average Degree | 2013 SSE | 2013 r |
|---|---|---|
| $k_C = 8; k_E = 6$ | 3.68 | 0.57 |
| $k_C = 9; k_E = 9$ | 3.57 | 0.554 |

#### Section 6: Estimating DMV per contact infectiousness τ, the rate of coastal and estuarine mixing α, and seasonal changes to {β}

6.1 Estimating τ

We inform the DMV per contact infectiousness τ:

$$\tau = \frac{R_0 \mu}{k}$$

Where the basic reproduction number of DMV ($R_0$) is 1.8[17], the recovery rate (μ) is equal to 1 over the DMV infectious period of 8 days[17], and the average synchrony degree ($k$) of bottlenose dolphins across ecotypes over the course of a day is 7 (established in supplement section 5.2). *We therefore calculate our standardized value of* τ *to be 0.032.*

6.2 Estimating α

At time of writing, the PCDP has completed 4 year sighting histories for all individuals sighted between 2015-2018 (n = 1,160 across 131 surveys). We use the same cluster methodology used in supplement section 5.1 of the main text on all 1,160 of these individuals to predict natural groupings according to their distance from shoreline. The elbow point, occurred at three clusters: a "nearshore" cluster (n=547 dolphins) with an average distance of 537m from shore (high confidence estuarine ecotype), a "midshore" cluster (n=373 dolphins) with an average distance of 1463m from shore (undetermined ecotype), and a "farshore" cluster (n=240 dolphins) with an average distance of 3605m from shore (high confidence coastal ecotype). We then dropped any survey in which dolphins of unconfirmed ecotype were present if high confidence dolphins of both ecotypes were not also present, leaving us with 47 surveys. The proportion of these 47 surveys in which both estuarine and coastal designated individuals were present was 6%. Since synchronized breathing is an affiliative behavior thought to occur between individuals with close bonds, we assumed that the rate of mixing between individuals of different ecotypes, α, would not exceed this value, *and thus determine* α *to range between 0-0.06.*

6.3 Estimating seasonal changes to β

We tested the effect of seasonal changes to transmission rate β due to behavioral and environmental factors. Recall that β at any time point in our the model can be calculated as:

$$\beta = k\tau$$

Where $k$ is average degree (number of synchrony contacts), and τ is the infectiousness of DMV from supplement section 6.1. We assume that seasonal changes to behavior will affect $k$ and seasonal changes to the environment will affect τ.

For the behavioral change, we assume that β is higher in the breeding season (between April 15 and July 7th, Figure S7) compared to the rest of the year due to a higher synchrony degree $k$ during this time period. Because the majority of our PCDP focal data on synchrony behavior (from supplement section 5) was collected during this time period, we assume that this data is reflective of the breeding season and therefore reduce $k$ outside of this time period. To determine by how much to reduce $k$, we refer to results from previous work[15] in which we found that individuals have high assortativity in their synchrony contacts. In other words, individuals preferentially sync with individuals of their same age and sex class and only 20-33% of synchrony contacts are outside of an individual's own age and sex class. Given this strong assortativity of synchrony behavior, we assume that these "outside" contacts would be the ones most likely to be removed outside of the breeding season, and thus reduce $k$ outside of this time period by 20% (the low end of this "outside" contact range).

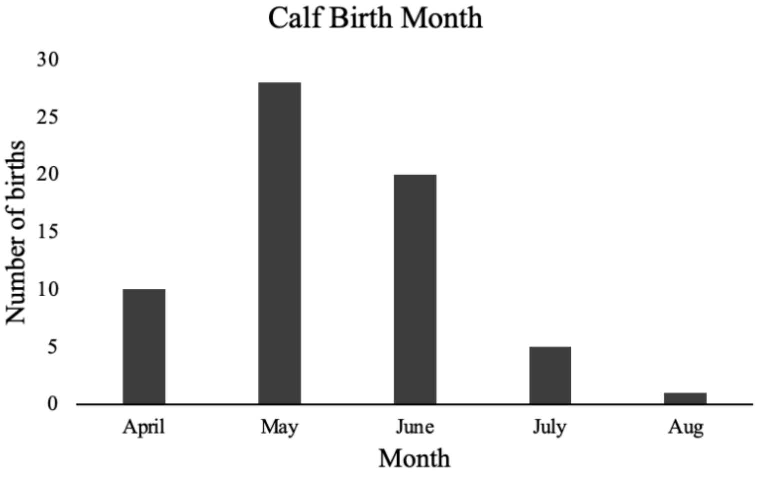


**Figure S6. Distribution of neonate birth dates by month between 2015-2019 at the PCDP field site (n=64).** Data and analysis courtesy of Jessica Wang, Ann-Marie Jacoby and Andrew Read in fulfillment of the Undergraduate Honors Thesis at Duke University.

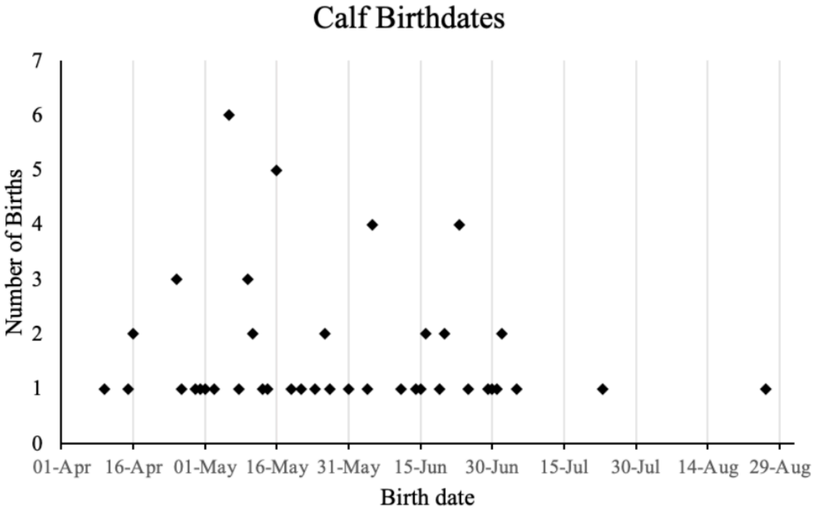


**Figure S7. Distribution of neonate birthdays by 15-day increments between 2015-2019 at the PCDP field site (n=64).** Data and analysis courtesy of Jessica Wang, Ann-Marie Jacoby and Andrew Read in fulfillment of the Undergraduate Honors Thesis at Duke University.

For the environmental change, we could not find a quantitative value on how much per contact infectiousness ($\tau$) might be increased in DMV due to colder temperatures. Therefore, we assume the same 20% reduction of $\tau$ outside of the cold water season (January 1-March 31) for a consistent comparison between the two seasonal scenarios.

6.4 Sensitivity analysis on seasonal changes to $\beta$

To assess how the magnitude of $k$ or $\tau$ reduction, and length of the breeding or cold water season affects our results, we examined additional models where these parameters are slightly adjusted (Table S5) to represent other potential **realistic** reduction and seasonal timing scenarios. For the seasonal social behavior hypothesis, we consider models with 1) a slightly shorter breeding season (May 1-June 30) representing the two months with the highest birthing peaks observed by the PCDP (Figure S6), and 2) a slightly longer breeding season (April 1 -July 30) to encompass the entirety of months with observed births. We also consider a loss of 10 and 35% of contacts outside of the breeding season (based on the percent range (20-33%) of outside demographic group contacts noted in 6.3[15]); since breathing synchrony contact is affiliative occurring between closely bonded individuals, we assume a further reduction of $k$ to be unrealistic. For the seasonal environment hypothesis, we consider a longer cold water season based on average sea surface temperatures (Figure S4) that encompasses November 1- May 31, and consider the same 10 and 35% reduction in $\tau$ for comparison.

We calculated the sum squared error (SSE) of each adjusted model's weekly infection results to the 2013 outbreak data, and use a one-way ANOVA and Tukey's HSD test to determine if these values of fit differ significantly from the values calculated from our initial seasonal behavior and seasonal environment models (Table S5).

*6.4.1 Results (Table S6):*

When reducing $\beta$ to reflect seasonal social behavior changes we see that altering the time of the breeding season to be slightly shorter or slightly longer than our defined breeding season of April 15-June 30 does not significantly change the infection time series of the model based on the SSE values calculated with the 2013 outbreak data. When adjusting the reduction of $k$, we also see no difference. We thus conclude that different, realistic, scenarios of seasonal behavior changes will not produce different results from our specified values.

When changing $\beta$ to reflect seasonal environmental changes, altering the time of the cold water season to be longer than our defined cold water season of January 1-March 31 does not significantly change the infection time series of the model based on the SSE values calculated with the 2013 outbreak data. Further reducing values of $\tau$ outside of the cold water season by 35% significantly increases these SSE values, indicating an even less consistent model. Reducing $\tau$ by 10% significantly decreases the SSE value, but this value is still higher than all values produced by any of the realistic seasonal behavior models. This indicates that no matter how $\beta$ is adjusted to account for higher infectivity in colder time periods, this hypothesis does not best represent past DMV outbreaks.

**Table S6. Sensitivity analysis on $k$ or τ reduction magnitude, and breeding or cold water season lengths for the seasonal social behavior and environment models.** Each row represents a model; the models with asterisks show the original parameters. All other rows show the adjustment value for either season time period or the magnitude of β reduction. Bold SSE values indicate that the associated adjusted model has a significantly different SSE value ($p<0.05$) to the 2013 outbreak data when compared to the SSE values of the original models (*). Lower SSE indicates a model that is more consistent with the 2013 outbreak data.

| Seasonal Hypothesis | Season time period | Reduction in $k$ or τ outside of season | 2013 SSE |
|---|---|---|---|
| Behavior* | Breeding: April 15-June 30 | $k$: 20% | 3.65 |
| Behavior | Breeding: April 1- July 31 | $k$: 20% | 3.39; p = 0.537 |
| Behavior | Breeding: May 1-June 30 | $k$: 20% | 3.67; p = 1.000 |
| Behavior | Breeding: April 15-June 30 | $k$: 10% | 3.83; p = 0.842 |
| Behavior | Breeding: April 15-June 30 | $k$: 35% | 3.78; p = 0.955 |
| Environmental* | Cold water: Jan 1-March 31 | τ: 20% | 4.68 |
| Environmental | Cold water: Nov 1- May 31 | τ: 20% | 4.46; p <0.533 |
| Environmental | Cold water: Jan 1-March 31 | τ: 10% | **8.29; p <0.001** |
| Environmental | Cold water: Jan 1-March 31 | τ: 35% | **3.99; p <0.001** |

#### Section 7: Epidemiological metapopulation model with stochastic and discrete integration of disease and movement dynamics

7.1 Disease dynamics

For ecotype $s$ in each patch $p$ for a compartment $X_s^p$, $X \in S, I, R$, we extracted a random variable using a binomial distribution for each possible transition out of that compartment in the discrete time interval $\Delta t$. This variable, $B_s^p(X_s^p, Y_s^p)$ is thus the number of dolphins that transition from $X_s^p$ to $Y_s^p$ in $\Delta t$.

For the transition from $S_s^p$ to $I_s^p$, in $\Delta t$ we sum two random variables $B_s^p(S_s^p, I_s^p)$ extracted from the binomial distribution:

$$Pr^{bin}(S_s^p,\ P_{S_s^p \to I_s^p})$$

The first variable is extracted with the transition probability:

$$P_{S^p \to I^p} = \lambda^p \Delta t$$

$\lambda_s^p$ is known at the force of infection for stock type $s$ for patch $p$ which is $\lambda_s^p = \beta_{ss} \frac{I_s^p}{N_s^p}$ where $\beta_{ss}$ is the transmission rate established for within ecotype $s$ transmission. $I_s^p$ is the number of infected dolphins in patch $p$ of ecotype $s$, and $N_s^p$ is the total number of dolphins in patch $p$ of ecotype $s$.

The second variable is extracted with the transition probability:

$$P_{S^p \to I^p} = \lambda_{mix}^p \Delta t$$

Where $\lambda_{mix}^p$ represents the force of infection for ecotype $s$ in patch $p$ coming from the other ecotype. $\lambda_{mix}^p = \beta_{CE} \frac{I_r^p}{I_r^p}$ where $\beta_{CE}$ is the transmission rates between ecotypes. $I_r^p$ is then number of infected dolphins in patch $p$ of the other ecotype $r$, and $N_r^p$ is the total number of dolphins in patch $p$ of the other ecotype $r$.

For the transition from $I^p$ to $R^p$, in $\Delta t$ we extracted a random variable $B_s^p(I_s^p, R_s^p)$ from the binomial distribution:

$$Pr^{bin}(I_s^p,\ P_{I_s^p \rightarrow R_s^p})$$

with the transition probabilities:

$$P_{I_s^p \rightarrow R_s^p} = \mu \Delta t$$

Where $\mu = 1/G$ and $G$ is the average infectious period of DMV, 8 days[17].

7.2 Movement dynamics

For coastal individuals, we assume that migration occurs between October-December and April-June. For estuarine individuals we assume that dispersal is year round, but northern dispersal (e.g., patch 2 to patch 1) occurs between January to June and southern dispersal (e.g., patch 1 to patch 2) occurs between July and December.

These yearly patterns have an integration time scale of 1 day. If $t$ falls during a migration or dispersal season in our simulation, we simulated the number of individuals of stock type $s$ in compartment $X_s^p$ traveling from patch $p$ to patch $q$ in $\Delta t$ as a random variable extracted from a binomial distribution:

$$Pr^{bin}(X_s^p,\ P_{X_s^p \rightarrow X_s^q})$$

With movement probabilities:

$$P_{X_s^p \rightarrow X_s^q} = \eta_{s,\,pq} \Delta t$$

Where $X_s^p$ is the number of dolphins of ecotype $s$ in compartment $X$ in patch $p$ at time $t$ and $\eta_{s,\,pq}$ is the movement rate for ecotype $s$ from patch $p$ to patch $q$.

#### Section 8: Establishing outbreak dataset from dolphin stranding data

We obtained data on bottlenose dolphin strandings (when a sick, injured or dead dolphin is found washed ashore or floating) between New Jersey and Georgia during the 2013 DMV outbreak from the National Oceanic and Atmospheric Administration (NOAA), and the 1987 DMV outbreak from the Smithsonian Division of Marine Mammal Collections. For both datasets we only considered mortalities that were not skeletal/mummified remains as the time of death for these can not be assumed to be during the associated epidemics.

According to NOAA, the peaks in strandings from the DMV epidemics spanned 21 months (1-Jul-13 to 1-Mar-15) and 11 months (1-Jun-87 to 31-May-88). Based on these dates we assume that the epidemics likely began in the middle of our model's migratory season 1 (April-June) and thus consider strandings that occurred beginning on 1-April (beginning of migratory season 1) until 31-Sep (end of the warm water season) of the following year to encompass a full migration cycle.

However, not all recorded strandings during this time period would be due to DMV. Therefore, in order to obtain a better idea of the true epidemic specific sickness and mortality, we removed the 'excess mortality' for each demographic group. We applied two generalized linear models to stranding data for 1) the three years prior to both epidemics, and 2) during the first year of each epidemic. The number of strandings per year was the response variable, with a categorical predictor variable for the latitude (rounded up to the nearest degree) at which the stranding occurred. For the pre-epidemic model we also include an interaction term for the year of the stranding. From our glms, we found that dolphins were observed stranded at an average rate of 0-14 strandings per year depending on latitude in the three years prior to the 1987 epidemic, and 2-50 strandings per year in the three years prior to the 2013 epidemic. During the 1987 epidemic, individuals stranded at a rate between 1-230 strandings per year, and between 49-372 strandings per year during the 2013 epidemic. Past work[17] assumed that the probability of a case recorded during a DMV outbreak being an actual DMV case is equal to the pre-epidemic stranding rate divided by the epidemic stranding rate for each latitude. Therefore, to remove excess mortality, we randomly removed a stranding from the data if it occurred during the epidemics, with the associated probability based on its latitude.

We take the resulting dataset and infer infection dates for each stranding based on the reported status of decay, and estimates of time from infection to death for morbilliviruses. Each stranding had an associated code between 1 and 4 that described its decay status, and we use estimates illustrated by a piglet from the Australian Museum[18] to determine a prospective death date: 1= Live animal (0 days since death), 2= Fresh dead (Initial decay; 0-3 days since death), 3= Moderate Decomposition (Putrification; 4-10 days since death), 4 = Advanced Decomposition (black putrification; 10-20 days since death). For each stranding we subtract a random number of days from its observation date drawn from the range associated with its decay status code to get an estimated death date. There is no data on the time from infection to death for DMV, but some studies suggest this time period varies between 2-3 weeks for canine[19] and phocine distemper[20] viruses, different strains of morbillivirus. Therefore, we subtract an additional number of days from the death date, randomly selected between 14 and 21 to get an estimated infection date. We refer to these final, adjusted, datasets with estimated infection dates as the "outbreak data".

### Section 9: Additional Results

#### 9.1 η calculations

**Table S7. The mean and standard deviations used to create distributions for the epidemiological metapopulation model movement rates.** At each time point in the model, a value for each parameter is drawn from the associated distributions to allow for variation in each parameter over the course of the epidemic.

| Parameter Symbol | Parameter Explanation | Mean | Standard deviation |
|---|---|---|---|
| $\eta_{C,12}$ | Coastal movement rate between patches 1 and 2 | 0.026 | 0.006 |
| $\eta_{C,23}$ | Coastal movement rate between patches 2 and 3 | 0.043 | 0.007 |
| $\eta_{C,34}$ | Coastal movement rate between patches 3 and 4 | 0.100 | 0.06 |
| $\eta_{E,12}$ | Estuarine movement rate between patches 1 and 2 | 0.034 | 0.002 |
| $\eta_{E,12}$ | Estuarine movement rate between patches 2 and 3 | 0.033 | 0.001 |
| $\eta_{E,12}$ | Estuarine movement rate between patches 3 and 4 | 0.041 | 0.002 |

9.2 Start Time and Patch Model Results

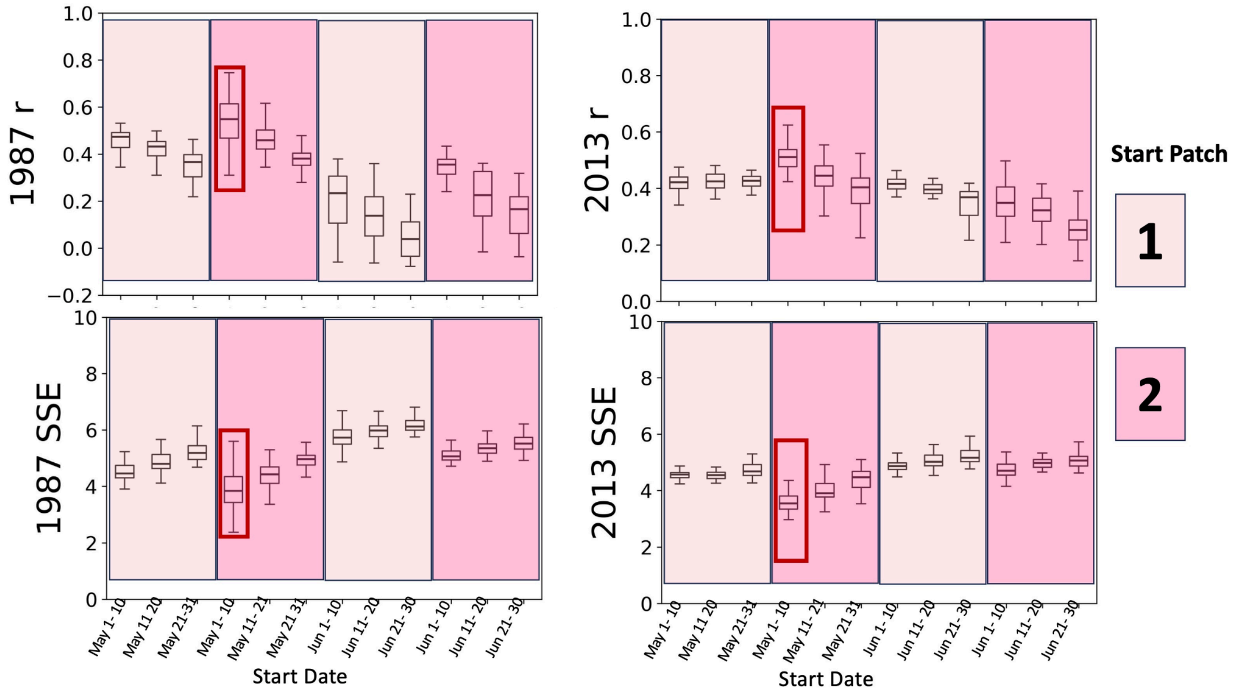


**Figure S8. Model fits for different start times and locations.** The Pearson's correlation coefficients (r), top, and sum squared error (SSE), bottom, for models with different infection start times between May and June and locations between patch 1 and 2. Higher r and lower SSE values are more consistent with the outbreak data for 1987 (left) and 2013 (right). We see that a model beginning infection between May 1-10 in patch 2 is most consistent with both the 2013 and 1987 outbreak data (red boxes) based on these values.

9.3 Seasonal Model Reuslts

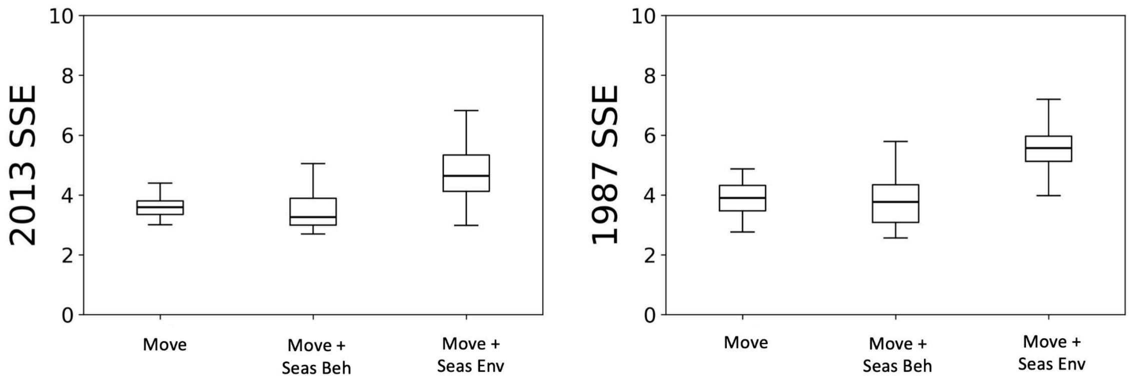


**Figure S9. The sum square error (SSE) values of each model calculated with the 2013 (left) and 1987 (right) outbreak data.** Lower SSE values indicate models that are more consistent with outbreak data. We see that a model with contact structured by seasonal behavior, is most consistent with DMV outbreak data from past epidemics based on these values.

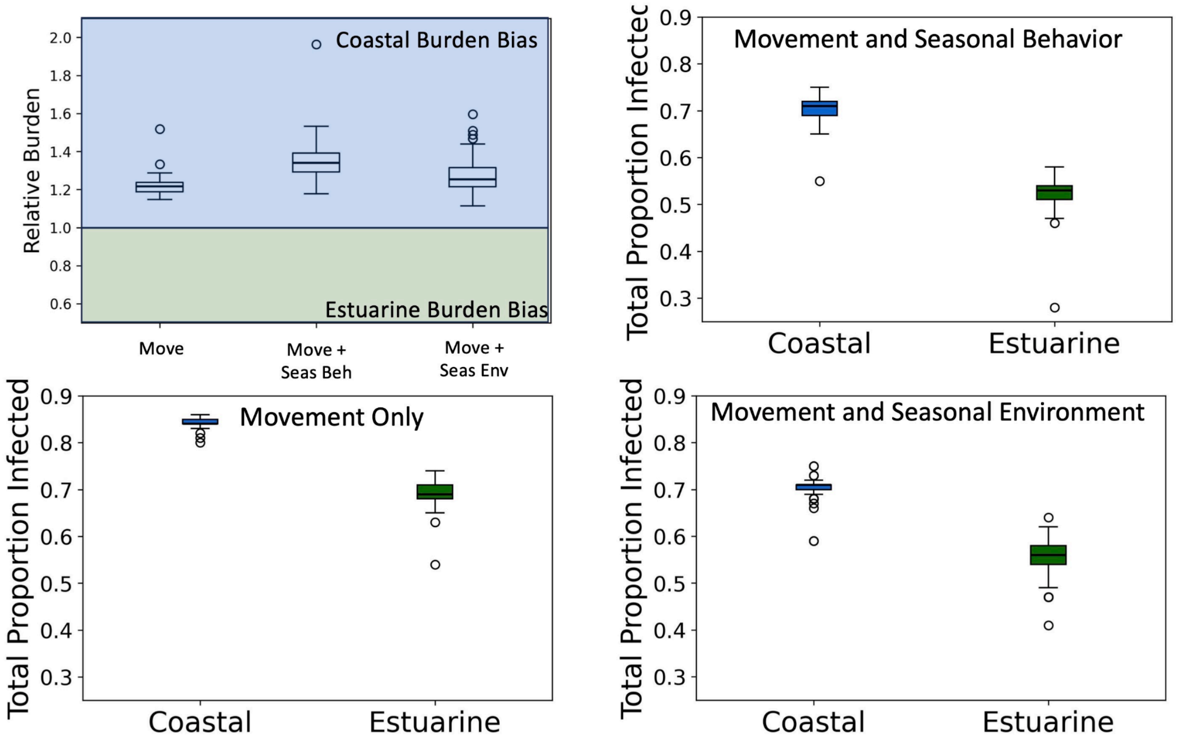


**Figure S10. The difference in coastal and estuarine infection burden for models that control for seasonal change.** Top left shows relative burden by ecotype, where relative burden is the ratio of the proportion of coastal individuals infected to estuarine individuals infected. When relative burden is equal to 1 (dotted line), there is no bias in infection burden between the two ecotypes. Values greater than 1 indicate a bias towards coastal infections, and values less than 1 indicate a bias towards estuarine infections. Remaining boxplots show the proportion of each ecotype infected in each model. Both seasonal change models maintain a higher coastal to estuarine infection burden observed in the movement only model, but the model that controls for seasonal behavior changes increases this value, providing more support for expert opinion.

#### 9.4 Control Model Results

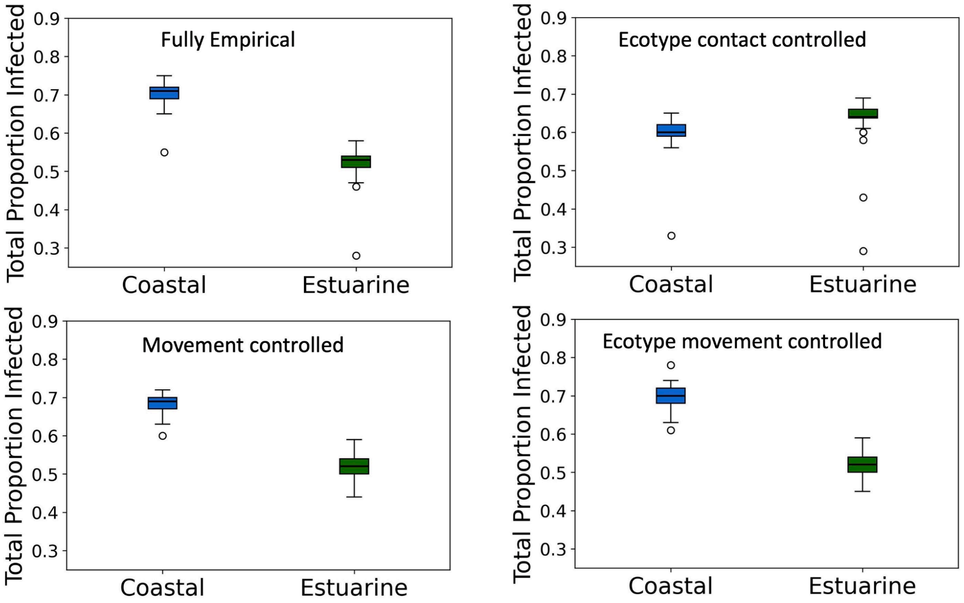


**Figure S11. The total proportion of each ecotype infected in each of the control models.**

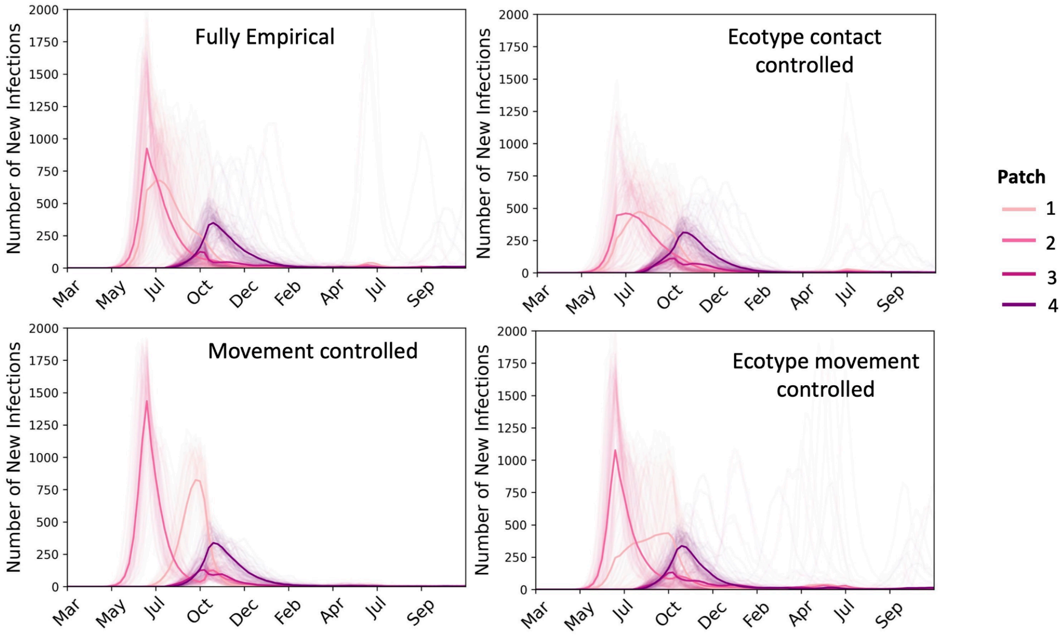


**Figure S12. The time series of weekly infections for three control models compared to the fully empirical model.**

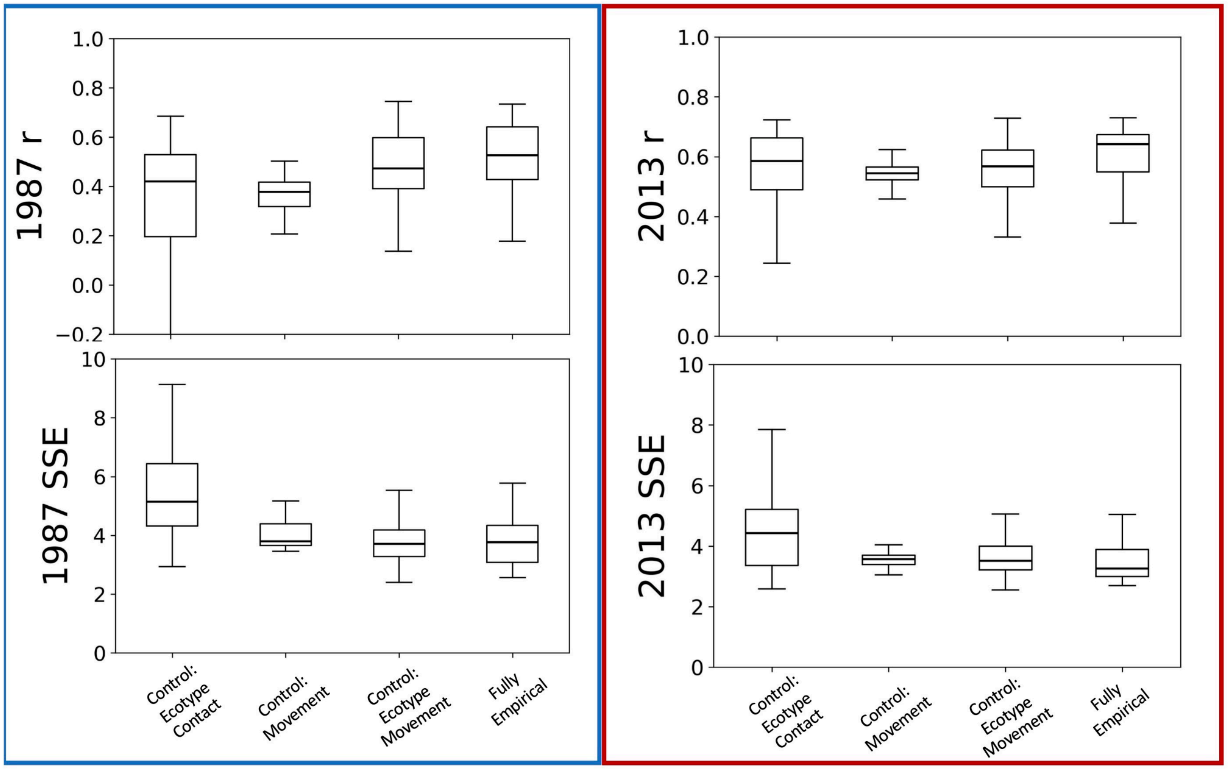


**Figure S13. Model fits for control models compared to the fully empirical model.** The Pearson's correlation coefficient (r), top, and sum squared error (SSE), bottom, for the control model infection results calculated with the 1987 (blue box) and 2013 (red box) outbreak data. Lower SSE and higher r values indicate models that are more consistent with the outbreak data.

9.5 Epidemic Likelihood and Surveillance Results

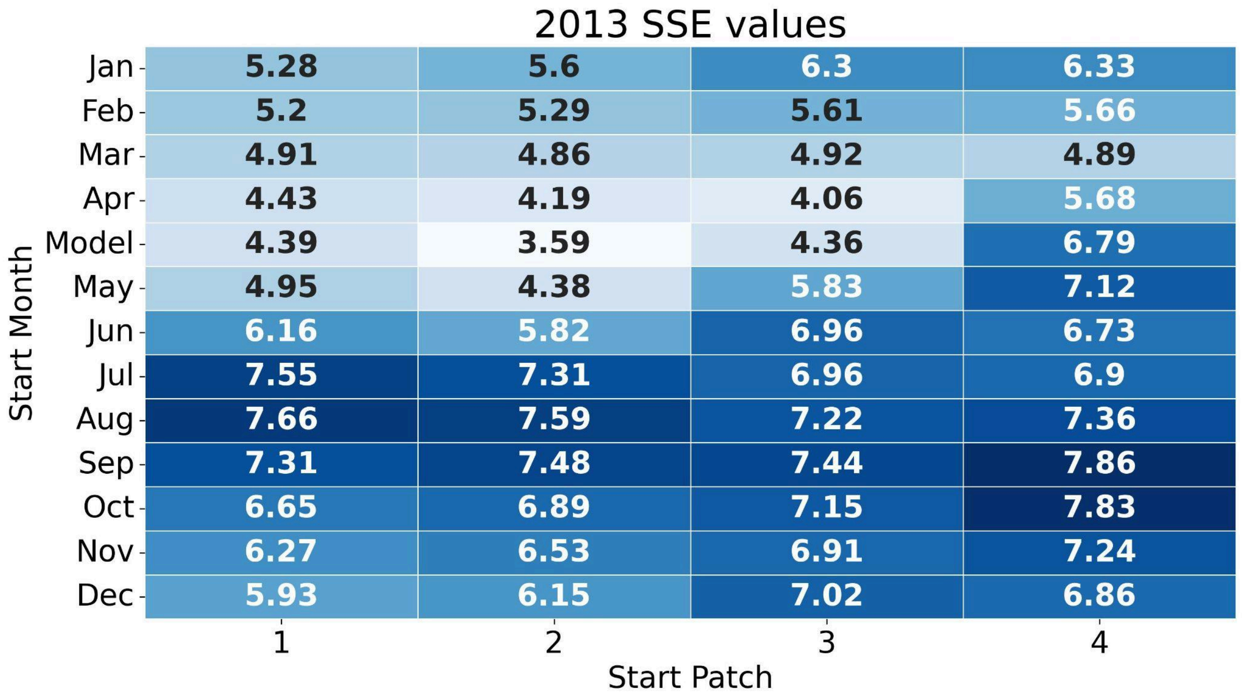


**Figure S14. The sum squared error (SSE) value for each possible start month and patch scenario calculated with the 2013 outbreak data**. Lower SSE values indicate a model is more consistent with the outbreak data.

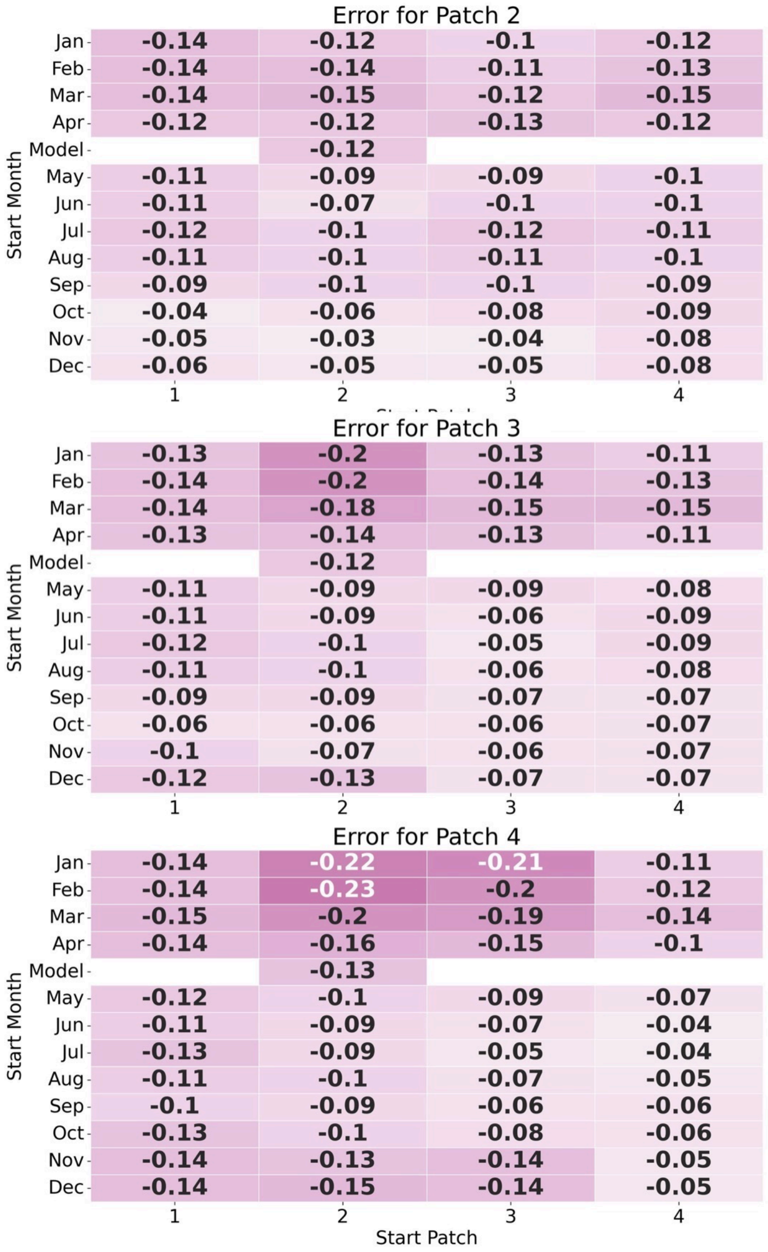


**Figure S15. The average difference in incidence for estuarine individuals in each patch and the entire coastline (error) for each start month and patch scenario.** Values closer to zero indicate estuarine individuals are better sentinels for the entire coastline for that scenario. The error for patch 2 is generally closest to 0 across all scenarios, and thus the best patch for surveillance.

#### Section 10: Considering the Florida coastline

10.1 Justification and Methods

One limitation of this work was that we do not consider the Florida coastline in our analysis due to the lack of consistent data present within the MABDC at this part of the coastline. There is also less known about the movements of stocks in the southern US Atlantic (e.g. the Southern migratory coastal, the South Carolina Georgia Coastal, the Northern Florida Coastal and the Central Florida Coastal) and whether or not they move into or out of Florida waters to mix with each other. However, the Florida coast did see a significantly high number of DMV related strandings during the 1987 and 2013 outbreaks.

Therefore, to inform metapopulation structure in this part of the coastline and better understand what movement to Florida is best supported by the observed disease dynamics, we produced counterfactuals using our epidemiological model. We created a theoretical fifth patch to encompass the northern Florida coastline from the GA/FL border to the border between Volusia and Brevard counties, since dolphins south of this part of the coastline were also involved in another mortality event caused by red tide in 2013. We populate this patch with 6000 coastal individuals and 1000 estuarine individuals based on the best estimates of stock sizes in this region prior to the 2013 outbreak[5]. We then test two possible hypotheses:

1. That coastal dolphins do not migrate into/out of Florida waters and that infection into Florida might have occurred from more sporadic, chance, interactions between individuals in patches 4 and 5.
2. Coastal dolphins migrate into and out of Florida waters seasonally at a rate of movement lower than those estimated between other patches (since the rate of migration into and out of Florida waters is unknown).

In the first hypothesis, we allowed for a sporadic infection to occur in patch 5 in September, the end of the warm water season, when coastal stocks are less likely to be delineated into distinct habitats and have the greatest potential for a sporadic interaction with individuals in patch 4. However, we set the rate of movement between patches 4 and 5 ($\eta_{45}$) to 0, which does not allow any individuals to migrate into and out of this patch.

In the second hypothesis, we set $\eta_{45}$ to be an average of 0.01 (standard deviation = 0.006) to represent a low rate of movement of coastal individuals to and from these two patches (based on our other $\eta$ estimates).

10.2 Results (Figure S16)

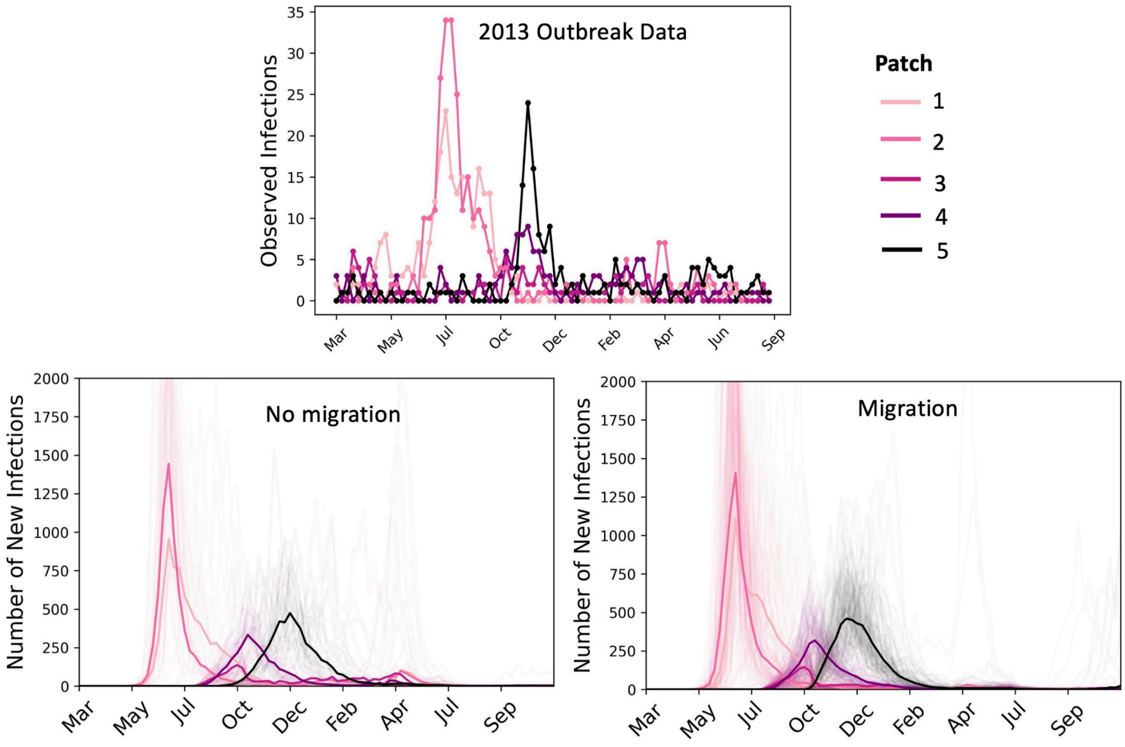


**Figure S16. The infection time series of a model that considers no migration into patch 5 (only sporadic interactions and subsequent infection) (bottom left), and migration into patch 5 (bottom right).** We see that both models are consistent with DMV outbreak data from the 2013 epidemic (top) based on the visual comparisons of the time series.

This analysis provides two potential avenues of infection in this part of the coastline. First, coastal individuals migrating into Florida from patch 4 (even at a slow rate of movement, $\eta_{C,45} = 0.01$) can reproduce infection dynamics similar to the 2013 DMV outbreak with a very high (97%) epidemic probability. Second, if coastal individuals do not migrate into Florida waters, sporadic interactions between coastal individuals in northern Florida and southern Georgia will also result in similar dynamics, but this scenario has a significantly lower probability of occurring (33%). These results suggest that migration likely does occur into Florida, and is the likely mechanism for infection in this region.

#### References


1. Thompson, J. W. *et al.* finFindR: Automated recognition and identification of marine mammal dorsal fins using residual convolutional neural networks. *Mar. Mammal Sci.* **38**, 139–150 (2022).
2. Urian, K. *et al.* Recommendations for photo-identification methods used in capture-recapture models with cetaceans. *Mar. Mammal Sci.* **31**, 298–321 (2015).
3. Hayes, S. A. *et al. U.S. Atlantic and Gulf of Mexico Marine Mammal Stock Assessments 2022*. *NOAA Tech Memo NMFS NE 304* (2023).
4. Waring, G. T., Josephson, E., Maze-Foley, K. & Rosel, P. E. *US Atlantic and Gulf of Mexico Marine Mammal Stock Assessments, 2013*.

https://repository.library.noaa.gov/view/noaa/4757 (2014).
5. Waring, G. T., Josephson, E., Maze-Foley, K. & Rosel, P. E. *US Atlantic and Gulf of Mexico Marine Mammal Stock Assessments 2015. NOAA Tech Memo NMFS NE 238* 501 (2016).
6. Balmer, B. *et al.* Ranging patterns, spatial overlap, and association with dolphin morbillivirus exposure in common bottlenose dolphins (*Tursiops truncatus*) along the Georgia, USA coast. *Ecol. Evol.* **8**, 12890–12904 (2018).
7. Copernicus Climate Change Service (C3S). Sea surface temperature daily data from 1981 to present derived from satellite observations. Copernicus Climate Change Service (C3S) Climate Data Store (CDS) https://doi.org/10.24381/cds.cf608234 (2019).
8. Rushing, C. S. An ecologist's introduction to continuous-time multi-state models for capture–recapture data. *J. Anim. Ecol.* **92**, 936–944 (2023).
9. McClintock, B. T. *et al.* Uncovering ecological state dynamics with hidden Markov models. *Ecol. Lett.* **23**, 1878–1903 (2020).
10. Michelot, T. & Blackwell, P. G. State-switching continuous-time correlated random walks. *Methods Ecol. Evol.* **10**, 637–649 (2019).
11. Albert, A. Estimating the Infinitesimal Generator of a Continuous Time, Finite State Markov Process. *Ann. Math. Stat.* **33**, 727–753 (1962).
12. Chesapeake Bay Shoreline High Resolution. Chesapeake Bay Program (2023).
13. Toth, J. L., Hohn, A. A., Able, K. W. & Gorgone, A. M. Defining bottlenose dolphin (*Tursiops truncatus*) stocks based on environmental, physical, and behavioral characteristics. *Mar. Mammal Sci.* **28**, 461–478 (2012).
14. Read, A. J. *et al. Stock Discrimination of Bottlenose Dolphins along the Outer Banks of North Carolina; Implications for the Risk of Entanglement in Coastal Gill Net Fisheries. Final Report North Carolina Sea Grant Bycatch Reduction Marine Mammal Project 10 DMM 01* (2013).
15. Collier, M. A. *et al.* Breathing in sync: how a social behavior structures respiratory epidemic risk in bottlenose dolphins. 2023.12.01.569646 Preprint at https://doi.org/10.1101/2023.12.01.569646 (2023).
16. Scott, G. P., Burn, D. M. & Hansen, L. J. The dolphin dieoff: long-term effects and recovery of the population. in *OCEANS '88. 'A Partnership of Marine Interests'. Proceedings* 819–823 (IEEE, Baltimore, MD, USA, 1988). doi:10.1109/OCEANS.1988.794905.
17. Morris, S. E. *et al.* Partially observed epidemics in wildlife hosts: modelling an outbreak of dolphin morbillivirus in the northwestern Atlantic, June 2013–2014. *J. R. Soc. Interface* **12**, (2015).
18. News, O. H. M.-W. 9am-9pm T.-S. 9am-5pm A. 1 W. S. S. N. 2010 A. P. +61 2 9320 6000 www australian museum C. © 2024 T. A. M. A. 85 407 224 698 V. M. Stages of decomposition. *The Australian Museum* https://australian.museum/learn/science/stages-of-decomposition/australian.museum/learn/science/stages-of-decomposition/.
19. Nagao, Y. *et al.* An Outbreak of Canine Distemper Virus in Tigers (*Panthera tigris*): Possible Transmission from Wild Animals to Zoo Animals. *J. Vet. Med. Sci.* **74**, 699–705 (2012).
20. Visser, I. K. *et al.* Vaccination of harbour seals (*Phoca vitulina*) against phocid distemper with two different inactivated canine distemper virus (CDV) vaccines. *Vaccine* **7**, 521–526 (1989).
